# Co-evolutionary Landscape at the Interface and Non-Interface Regions of Protein-Protein Interaction Complexes

**DOI:** 10.1101/2021.01.09.426046

**Authors:** Ishita Mukherjee, Saikat Chakrabarti

## Abstract

Proteins involved in interactions throughout the course of evolution tend to co-evolve and compensatory or coordinated changes may occur in interacting proteins to maintain or refine interactions between them. However, certain residue pair alterations may prove to be detrimental for functional interactions. Hence, determining co-evolutionary pairings that could be structurally or functionally relevant for maintaining the conservation of an inter-protein interaction is important. Inter-protein co-evolution analysis in a number of complexes with the help of multiple existing methodologies suggested that co-evolutionary pairings can occur in spatially proximal as well as distant regions of inter-protein interaction complexes. Subsequently, the Co-Var (**Co**rrelated **Var**iation) method based on mutual information and Bhattacharyya coefficient was developed, validated and found to perform relatively better than CAPS and EV-complex. Interestingly, while applying the Co-Var measure on a set of protein-protein interaction complexes, co-evolutionary pairings were obtained in spatially proximal and distant regions in inter-protein complexes. Our approach involves determining high degree co-evolutionary pairings which include multiple co-evolutionary connections between particular co-evolved residue positions in one protein and particular residue positions in the binding partner. Detailed analyses of high degree co-evolutionary pairings in protein-protein complexes involved in inter-cellular communication during cancer metastasis were performed. These analyses suggested that most of the residue positions involved in such co-evolutionary pairings mainly occurred within functional domains of constituent proteins and substitution mutations were also common among these co-evolved positions. The physiological relevance of these predictions suggests that Co-Var can predict residues that could be crucial for preserving functional protein-protein interactions. Finally, **Co-Var** web server that implements this methodology was developed. This web server available at http://www.hpppi.iicb.res.in/ishi/covar/index.html identifies co-evolutionary pairings in intra-protein and inter-protein complexes.

## Introduction

Intra-protein co-evolution which involves compensatory substitutions within proteins can restore functionality by sustaining the fitness of the protein under constraints imposed by physico-chemical interaction forces, structural and folding associated factors (Fitch 1971; Chakrabarti and Panchenko 2010). Multiple approaches have been utilised to study intra-molecular co-evolution such as substitution pattern correlations, mutual information of amino acid frequencies between positions in a multiple sequence alignment (MSA), analysis of evolutionary phylogenetic trees etc. (deJuan et al. 2013). Further, a number of coevolution-based contact prediction methods which include adjustments for direct and indirect couplings have been developed for monomeric proteins (Marks et al., 2011; Morcos et al., 2011; Hopf et al., 2012; Jones et al., 2012; Kamisetty et al., 2013). In general, it has been observed that interacting residues in close proximity have a tendency to co-evolve (Pollock et al., 1999; Valencia and Pazos 2002; Choi et al., 2005; Chakrabarti and Panchenko 2009). However, large number of coevolutionary connections may occur at less variable positions within a protein family (Mandloi and Chakrabarti, 2017). Moreover, clusters of positions which are usually not in contact but tend to be located near binding regions or active sites have been found to co-evolve (Gloor et al., 2005).

Protein-protein interactions are inherently important in signal transduction pathways or metabolic reactions within cells to carry out diverse physiological processes. Single protein molecules may interact fleetingly in transient complexes involved in different types of interactions (for e.g. signalling–effector, enzyme–inhibitor, enzyme–substrate, hormone–receptor etc.) whereas some proteins may exist as parts of multi-subunit enzymes in permanent obligate interactions (Mintseris and Weng, 2005). During such interactions, correlative sequence evolution can occur between proteins that physically interact or have a functional association in a manner such that amino acid changes at one site in a molecule may give rise to changes in selection pressure at another site in the binding partner. This evolutionary interaction between protein sites in different molecules that undergo compensatory changes to maintain the stability or functions of the interaction over the course of evolution is referred to as inter-protein co-evolution (Lovell and Robertson, 2010). The observation that interface positions exhibit changes in a correlated manner among interacting molecules lead to the development of methods for predicting contacting pairs of residues from sequence information (Pazos et al., 1997). Analysis of co-evolution in inter-protein complexes has demonstrated that residues at the interfaces of obligate complexes co-evolve with their interacting partners whereas transient protein interaction complexes have an increased rate of substitution at the interface residues with less correlated mutations occurring across the interface (Mintseris and Weng, 2005). In general, while functionally important co-evolving residues having high mutual information (MI) occur in structural proximity (Marino Buslje et al., 2010; Teppa et al., 2017), a fraction of coevolving residue pairs predicted based on direct couplings have recently been shown to occur at distant regions between protein structures (Anishchenko et al., 2017).

Evolutionary pressure is likely to maintain an interaction between protein interfaces wherein selection restricts amino acid replacements or preserves a degree of conservation in the binding interfaces to maintain functionality of such interactions (Lovell and Robertson, 2010). In this respect, analysis of molecular co-evolution in inter-protein complexes may be useful for determining co-evolutionary pairings among interface residues and it is likely that co-ordinated changes at these residue positions are likely to be crucial for a functional interaction between these sets of proteins. In this study, we have developed a method named Co-Var (**Co**rrelated **Var**iation) aiming to determine both inter-protein and intra-protein residues that are likely to carry out crucial structural or functional roles in protein-protein interactions via establishing co-evolutionary pairings between themselves. Herein, we have determined the applicability of the Co-Var measure in studying inter-protein co-evolution and compared the methodology with selected protein-protein co-evolution analysis methodologies such as CAPS (Fares and McNally 2006; Fares and Travers 2006) and EV-complex (Hopf et al., 2014). However, we observed that co-evolving residues in inter-protein interaction complexes were found to occur in close spatial proximity in interface regions as well as in non-interface regions, an observation concordant with a previous study (Anishchenko et al., 2017). Based on this observation, we have considered the hypothesis that co-evolutionary pairings that occur in interface and non-interface regions could be crucial for native interactions and absence of co-ordinated changes at these positions are likely to contribute to altered interaction profiles and aberrant complex functionality. Therefore, to study the likely structural or functional relationship between co-evolutionary pairings in interface and non-interface regions and aberrant complex functionality certain protein-protein interaction complexes that exhibit frequent mutations in cancer have been considered. Studies on distribution pattern of disease associated variants have identified that disease causing mis-sense mutations frequently occur at the core region of protein-protein interaction interfaces or at the ligand-binding sites and residues involved in enzymatic function (Gao et al., 2015; David and Sternberg, 2015). Thus, co-evolution in certain ligand-receptor proteins as case studies has been studied with the help of Co-Var to exemplify the physiological relevance of predicted co-evolutionary pairings. Based on these analyses, we could determine that lack of co-ordinated changes at co-varying residue positions could be a likely contributing factor to the altered functionality of disease associated complexes involved in processes such as cancer metastasis. In this manner, one can ascertain co-evolutionary pairings that are likely to be crucial for functional interactions between proteins which when altered could be disease associated. Therefore, in this work we have implemented the Co-Var methodology, determined its applicability in studying inter-protein co-evolution and utilised it to study inter-cellular interaction complexes involved in cancer metastasis. The composite co-evolutionary measures along with various options to visualize such co-evolutionary pairings onto the representative structure of the complexes have been implemented within Co-Var web server which is freely accessible at http://www.hpppi.iicb.res.in/ishi/covar/index.html.

## Materials and Methods

### Determining co-evolutionary pairs in positive and negative protein-protein interaction complexes

A set of protein-protein interaction complexes (100) were collected from previous published data (Sowmya et al., 2015; Mintseris and Weng, 2003; Rodriguez-Rivas et al., 2016) and complexes satisfying our selection criteria of sufficient number of homologs, availability of crystal structure etc. were considered during this analysis. From this set, 50 protein complexes were selected as the set of interacting complexes that are likely to co-evolve (“positive set”) (Supplementary Table 1). Additionally, proteins which were known to be non-interacting based on experimental analysis were randomly selected from the Negatome database (Smialowski et al., 2009) as the “negative set” (Supplementary Table 1). The compiled positive and negative set comprised of 50 heterodimeric protein pairs each were subjected to MirrorTree (Ochoa and Pazos, 2010), CAPS (Fares and McNally 2006; Fares and Travers 2006) and EV-complex (Hopf et al., 2014) methods respectively. Other methods such as DCA or GREMLIN which are mainly used for contact prediction have not been considered here since we wanted to capture co-evolved positions in interface and non-interface regions to determine the possible implications of the same. Close orthologs or similar sequences were determined using DELTA-BLAST (Domain enhanced lookup time accelerated BLAST) (Boratyn et al., 2012) and taxonomy filtered non-redundant sequences having E-value <= 1E-04, query coverage >= 70%, sequence identity >= 45% were utilized for preparing multiple sequence alignments (MSA) representative of each sequence family with the help of MAFFT (Katoh et al., 2002). MirrorTree was utilized to determine whether the proteins co-evolve considering alignments of homologous sequences of the representative interacting and non-interacting proteins in each set. Here, tree similarities are quantified with the help of linear correlation by extracting inter-ortholog distance matrices from the MSA-derived trees of orthologous protein sequences in the two families (Ochoa and Pazos, 2010). CAPS was run with the help of default parameters on the set of alignments generated to identify amino acid co-variation with the help of BLOSUM corrected amino acid distances and phylogenetic sequence relationships (Fares and McNally 2006). EV-complex was utilized to predict inter-evolutionary couplings in the alignment of concatenated sequences (generated internally during the calculation) using a global probability model of sequence co-evolution (pseudo-likelihood maximization) (Hopf et al., 2014). Subsequently, a distance distribution plot was prepared to analyse the inter-residue distances between the predicted co-evolving residue positions among the interacting proteins (positive set) obtained from CAPS and EV-complex.

### Identifying intra-protein and inter-protein co-evolution with an information theory based measure (Co-Var)

In information theory, mutual information represents the entropy-based formulation for quantifying the interdependence between the values of two random categorical variables which in this case could be position-wise amino acid frequency distributions (de juan et al, 2013). Further, mutual information is defined as the amount of information one variable or amino acid frequency distribution (column A) can tell us about the value of another variable or amino acid frequency distribution (column B). In other words, it is the reduction in uncertainty (entropy) in the value of ‘A’, if we know the value of ‘B’ and is calculated considering the sum over all the possible combinations of di-residue frequencies (Dunn et al., 2008). Mutual information (MI) between two aligned columns A and B is calculated as

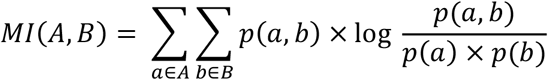

wherein, ‘p(a)’ is the frequency of occurrence of each residue in the column ‘A’, ‘p(b)’ is the frequency of occurrence of each residue in the column ‘B’ and ‘p(a,b)’ represents the di-residue frequency. Additionally, the Bhattacharyya coefficient quantifies the overlap between set of amino acids between a pair of columns. It is a measure of similarity between two datasets or distributions and is used to calculate the amount of overlap between two distributions, by splitting the samples into several partitions.

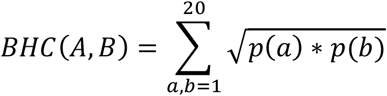

wherein, BHC(A,B) denotes Bhattacharyya coefficient between positions A and B, ‘p(a)’ and ‘p(b)’ are the amino acid residue frequencies present in the respective positions.

A score is computed considering homologous set of sequences within a protein family to derive intra-residue correlations in a single protein wherein each alignment position is compared to all other positions within the alignment for the protein family under consideration. Alternately, correlated positions between pairs of interacting proteins can also be identified based on the Co-Var score considering the position-wise amino acid frequencies in the multiple sequence alignments of the proteins involved in inter-protein interaction. Correlations between evolutionary patterns within proteins or between proteins may be determined based on the Co-Var score as outlined below:

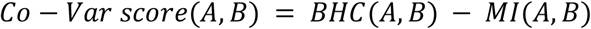

wherein, Co-Var score(A,B) represents the co-variation score between position A and B, MI (A,B) and BHC(A,B) denote mutual information and Bhattacharyya coefficient between positions A and B. Additionally, in case of intra-protein co-evolution ‘A’ and ‘B’ represent different positions within the multiple sequence alignment of a protein family whereas in inter-protein co-evolution analysis ‘A’ and ‘B’ represent a position in the alignment for the first protein family and another position in the second protein family respectively. The Co-Var methodology to study intra-protein and inter-protein co-evolution has been depicted in **Figure 1 and Figure 2**, respectively.

**Figure 1:**
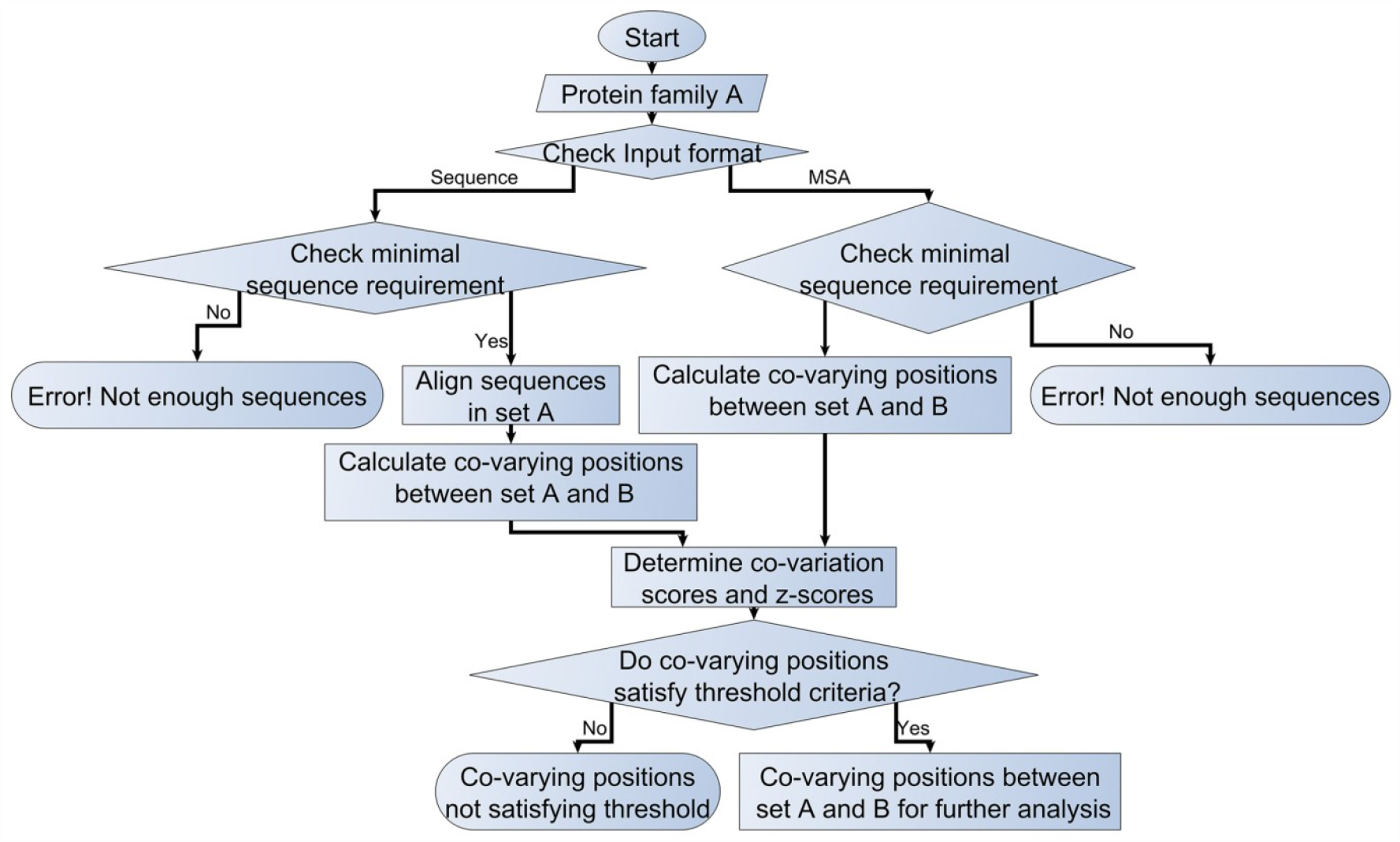
Co-Var methodology for studying intra-molecular co-variation in proteins.

**Figure 2:**
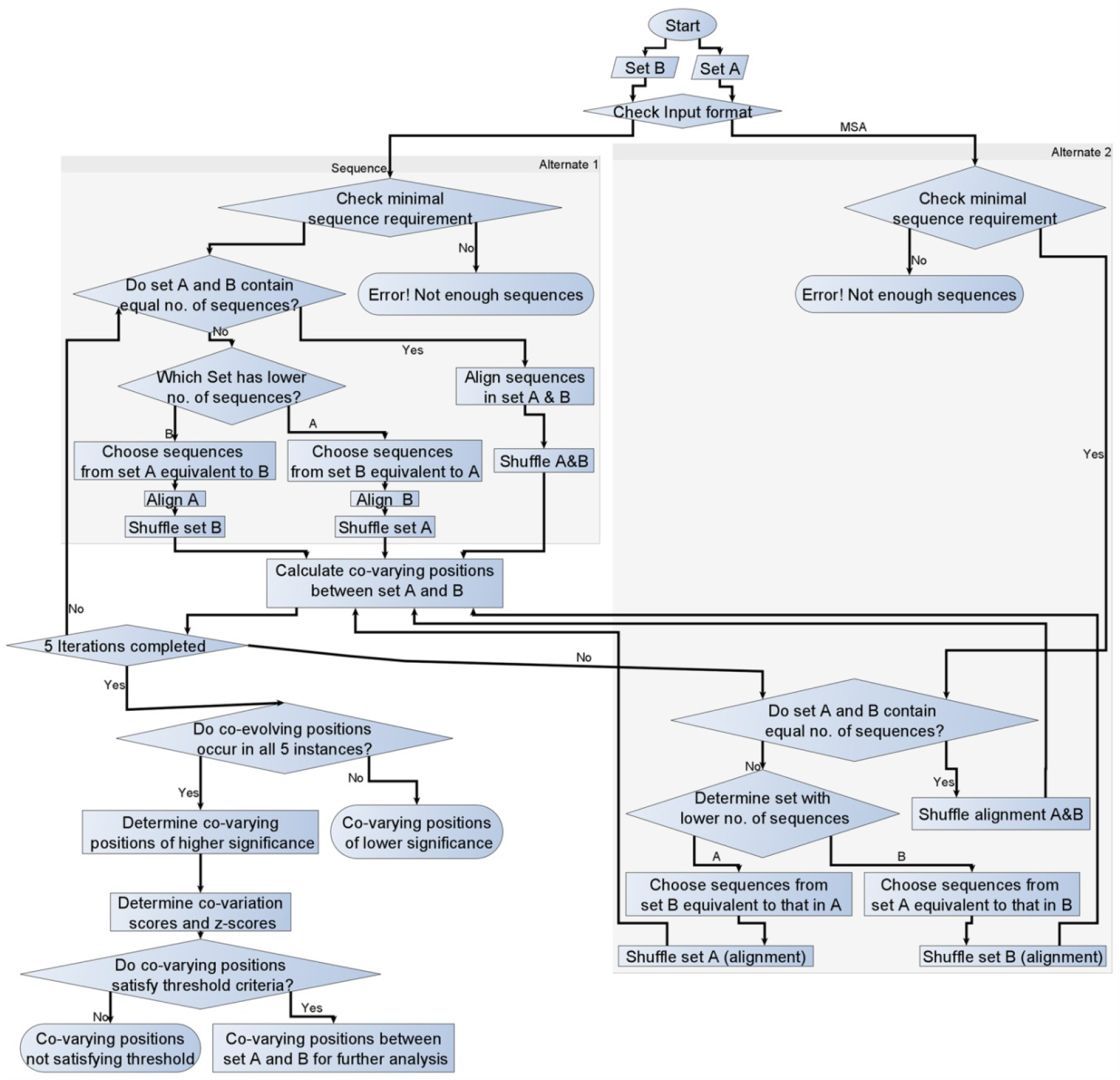
Co-Var methodology for studying inter-molecular or protein-protein co-evolution.

Close orthologs or similar sequences were determined using DELTA-BLAST (Domain enhanced lookup time accelerated BLAST) (Boratyn et al., 2012) and minimum 50 sequences were considered for generating an MSA representative of each family wherein the first sequence in each alignment was considered as the reference sequence. While detecting co-evolution, alignment shuffling was performed with a view to reduce the influence of phylogenetic relationships. Shuffling was performed by randomly selecting orthologous sequences in each family such that amino acids across the column exhibited variation. Further, multiple instances of the program were run and co-evolutionary pairings that are consistently identified across the different runs based on z-score threshold were considered such that additional statistical significance can be assigned to the co-evolving positions. After the calculation of the Co-Var scores, the residue pairs and scores were mapped to a corresponding reference sequence and structure for each set and average Co-Var score and corresponding z-scores across the runs were determined. Co-evolving positions having significant Co-Var score and z-scores which are reported in multiple runs (5) of the program with alignment shuffling have been considered for further analysis. Co-evolved positions were selected based on a z-score threshold where z-scores were calculated on the Co-Var score. Further, Co-Var scores corresponding to lower (negative) z-scores are indicative of higher likelihood of co-variation.

### Benchmarking of predicted co-evolutionary positions in protein-protein interaction complexes

In order to determine the applicability of Co-Var methodology in studying inter-protein co-evolution the ‘percentage of co-evolved pairs’ in interacting (positive) and non-interacting proteins (negative) has been considered as an index. Moreover, the ‘percentage of co-evolved pairs’ predicted for each positive and negative set pair by CAPS and EV-complex have been determined for these complexes. Based on these analyses, we have determined whether these indices can successfully differentiate between the positive and negative set. Subsequently, by considering a reference structure for each of the positive set of complexes, a distance distribution plot was prepared to analyse the inter-residue distances between the co-evolving residues predicted with the help of Co-Var. Additionally, two measures that capture the co-evolving pairs in the overall complex [Percentage of co-evolved pair that occur at interface (IC)] and those that occur at the interface [Percentage of interacting pair that are co-evolved (PC)] were also computed. For this purpose, co-evolving residues in inter-protein interaction complexes in close spatial proximity (<7Å) were considered as interface residues and other residues (inter-residue distances>7Å) were considered as non-interface residues.

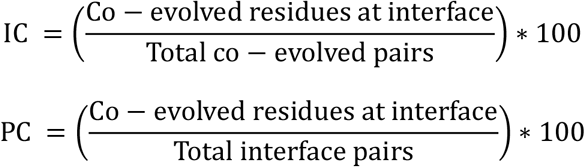

### Studying intra-protein co-evolution using the Co-Var methodology

A set of 252 conserved domain database (CDD) protein family alignments (Marchler-Bauer et al., 2015) with at least 80 sequences in each alignment were collected to study intra-molecular co-evolution (Supplementary Table 2). Intra-molecular co-evolution in these protein families was studied with the help of Co-Var, CAPS (Fares and Travers 2006), MI (Korber et al., 1993) and PSICOV (Jones et al., 2012) respectively. Methods were run considering default optimal parameters and the inter-residue distances among intra-protein co-evolved pairs predicted utilising these programs were determined for comparison.

### Studying co-evolution in intercellular protein-protein interaction complexes

In this study, inter-protein co-evolution has been studied in intercellular protein interaction complexes such as CSF3-CSF3R, TGFA-EGFR, FGF1-FGFR1, TGFB3-TGFBR2, FGF10-FGFR2 and FGF1-FGFR2. The respective sequences and structures were obtained from the UniProt and PDB databases (Rose et al., 2015; Tamada et al., 2006; Breuza et al., 2016; The UniProt Consortium, 2019). Orthologs in each family were determined with the help of DELTA-BLAST (Boratyn et al., 2012) and taxonomy filtered non-redundant sequences having E-value <= 1E-04, query coverage >= 70%, sequence identity >= 45% were utilized for preparing multiple sequence alignments in MAFFT (Katoh et al., 2002). Co-evolving residue positions among the representative sequences in each sequence family involved in the interaction complex were determined with the help of Co-Var methodology considering these alignments as input and z-score <= −1 as the selection threshold. Subsequently, co-evolved residue pair positions were mapped onto the corresponding 3D structure of the reference sequence to determine the inter-residue distances for a distance distribution analysis. The degree of each residue position among the co-evolved pairs was determined by analyzing a network representing residue positions as nodes and co-evolutionary pairings between positions as edges. Based on whether a residue position had a degree higher than the median of the degree distribution, high degree co-evolved positions were selected for further analysis. Moreover, in order to determine whether the high degree co-evolved positions are biologically relevant or not, a number of additional analyses were performed. It was studied whether these positions are present within the functional domain regions of each protein, whether these positions are important from the stand point of intra-molecular co-evolution or whether these positions are frequently prone to mis-sense substitutions.

In addition, the Co-Var methodology was also utilised to study co-variation within each protein involved in the inter-cellular protein interaction complex case studies considered herein. This analysis was performed in order to determine whether the high degree co-evolved residue positions identified in inter-molecular co-evolution could additionally have crucial roles within each protein in terms of their role in intra-molecular co-evolution.

### Investigating the functional and physiological relevance of the co-evolved sites

It would be interesting to ascertain whether the high degree co-evolutionary pairings have a biological significance. High degree co-evolved positions that are likely to be important for maintaining inter-protein functional interaction were determined and represented on a Circosplot (R core, 2017) for easy interpretation. The functional relevance of the predicted co-evolutionary pairings has been studied by performing domain mapping and mutation analysis. Details regarding the domains in each protein were obtained by querying the Pfam database (Finn et al., 2014) and in-house perl programs were utilized to determine whether the residue positions involved in co-evolutionary pairings occurred within the functional domain regions of the interacting proteins. Additionally, mutation data from the COSMIC database (Tate et al., 2019) has been considered to map whether residue positions important from the standpoint of inter-molecular co-evolution exhibit frequent substitution mutations in disease conditions such as cancer. As a result of the mutation it is plausible that the interaction between these proteins is perhaps compromised in conditions such as cancer. Amino acid pairing frequencies among the predicted co-evolved positions in the native reference sequences were compared to the ones obtained based on the assumption that the sequences exhibit the substitution mutations.

### Co-Var web server for predicting intra- and inter molecular co-evolutionary pairings

In order to provide a wider scope to the Co-Var methodology and make it applicable to a variety of biomolecules and bio-molecular complexes, we have generated a web server version of the method which is freely accessible to the users. Utilizing a set of homologous sequences or alignment(s) of proteins as inputs the Co-Var methodology may be utilized for studying intra-protein or inter-protein co-evolution analysis in our Co-Var web server available via http://www.hpppi.iicb.res.in/ishi/covar/index.html. The front end of the server is HTML, PHP and java based while a perl based implementation of the Co-Var methodology works on the backend of the server to predict reference sequence (first sequence in the alignment) mapped co-evolved residue positions. Further, based on an uploaded reference structure inter-residue distances between the co-evolutionary pairings and structural mapping of pairings can be obtained. Co-evolutionary pairings in close structural proximity can be visualized in a viewer (Rego and Koes, 2015). Additional modules are available for functional interpretation of the inter-protein co-evolutionary pairings in terms of their frequency of occurrence among predicted co-evolved positions (high degree co-evolved positions). Moreover, the list of reference sequence or structure mapped or functionally relevant co-evolutionary pairings can be downloaded from the results link.

## Results

### Studying co-evolution in protein-protein interaction complexes utilising different inter-protein co-evolution analysis methods

The positive and negative set of complexes were analysed in inter-protein co-evolution analysis programs [MirrorTree (Ochoa and Pazos, 2010), CAPS (Fares and McNally 2006; Fares and Travers 2006) and EV-complex (Hopf et al., 2014)] for comparison. MirrorTree method provides a correlation coefficient as an estimation of the likelihood of co-evolution between two protein families/alignments. Higher the correlation coefficient value, more likely it is that the proteins are co-evolving. However, it must be noted that MirrorTree does not provide residue wise coevolutionary information between a pair of proteins. For our positive dataset the median of the distribution for the MirrorTree correlation co-efficient was higher than 0.8 indicating that these complexes are more likely to co-evolve than the negative set of complexes that had the median of correlation co-efficient distribution lower than 0.8 (p-value <= 0.0001) (Figure 3A). Inter-residue distances were determined among these predicted co-evolved positions in the representative 3D structures. It was identified that a large percentage of co-evolved pairs did not lie in close spatial proximity (<10Å) in EV-complex and CAPS respectively (Figure 3B,C).

**Figure 3:**
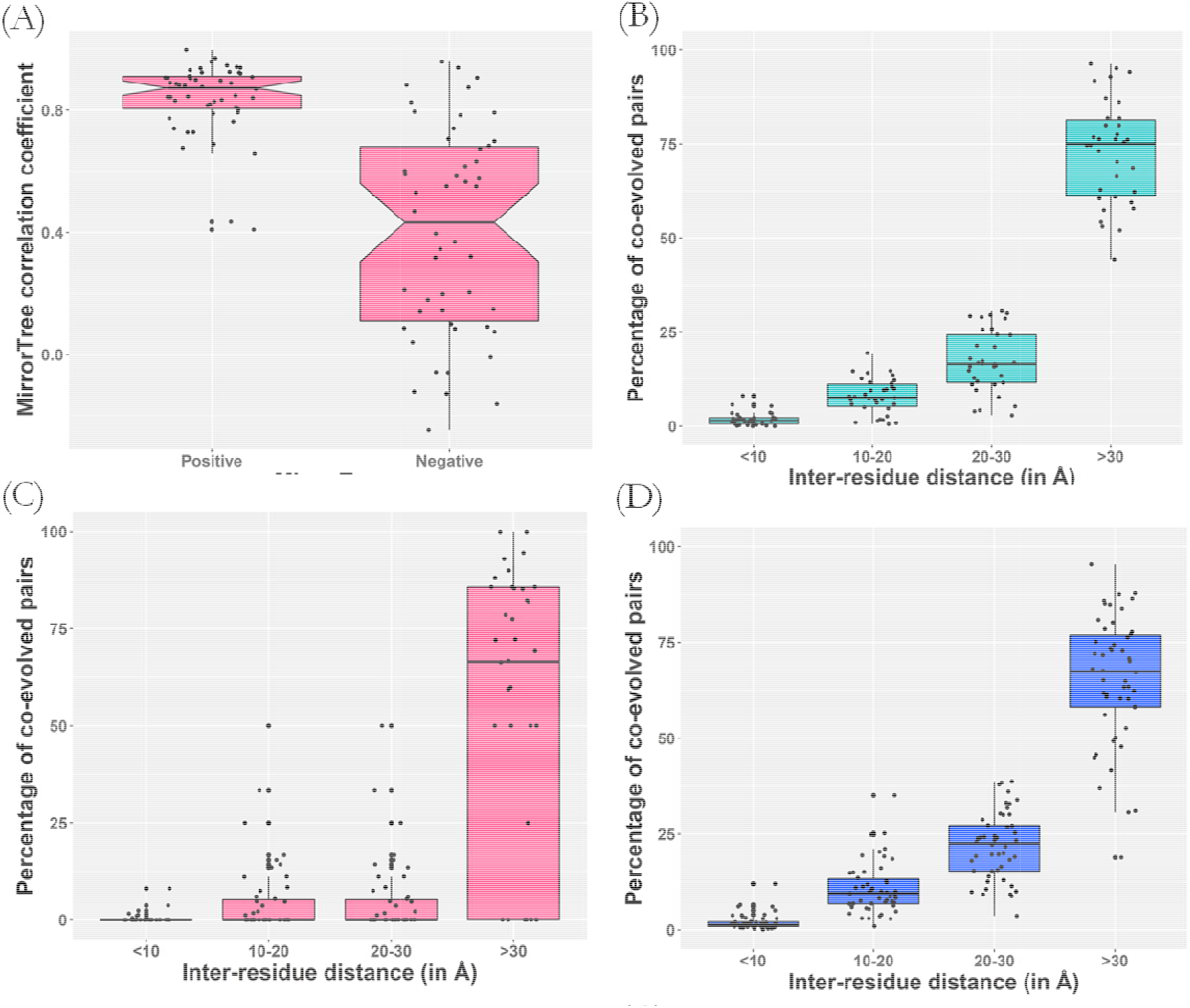
Analysis of inter-protein co-evolutionary pairings from a structural perspective. A set of interacting proteins which are likely to co-evolve (Positive) and a set of non-interacting proteins (Negative) that are less likely to co-evolve were studied with the help of multiple programs available to study inter-protein co-evolution. (A) MirrorTree based prediction of co-evolving and non co-evolving proteins considering the positive and negative dataset (B) Inter-residue distance distribution for EVcomplex predicted co-evolved pairs (C) Distance distribution analysis for CAPS based prediction of co-evolved pairs in interacting complexes (D) Distribution of ‘percentage of co-evolved pairs’ predicted by Co-Var in specific inter-residue distance distribution bins.

An evolutionary approach based on information theory may be utilised to identify inter-dependent protein residue positions that are crucial for conservation of interaction between two proteins. A significant fraction of residue pair positions were identified as co-evolving in each of the interacting protein complexes considered herein based on the Co-Var methodology (Supplementary Table 1). For this purpose, ‘percentage of co-evolved pairs’ has been determined among interacting (positive set [50 complexes]) and non-interacting proteins (negative set [50 complexes]) and this parameter is likely to be higher for interacting complexes which are likely to co-evolve compared to the non-interacting proteins which are less likely to co-evolve. In general, interacting complexes had a higher percentage of co-evolving pairs than the non-interacting complexes as suggested by inter-protein co-evolution analysis programs such as CAPS and EV-complex (Figure 4A, B).

**Figure 4:**
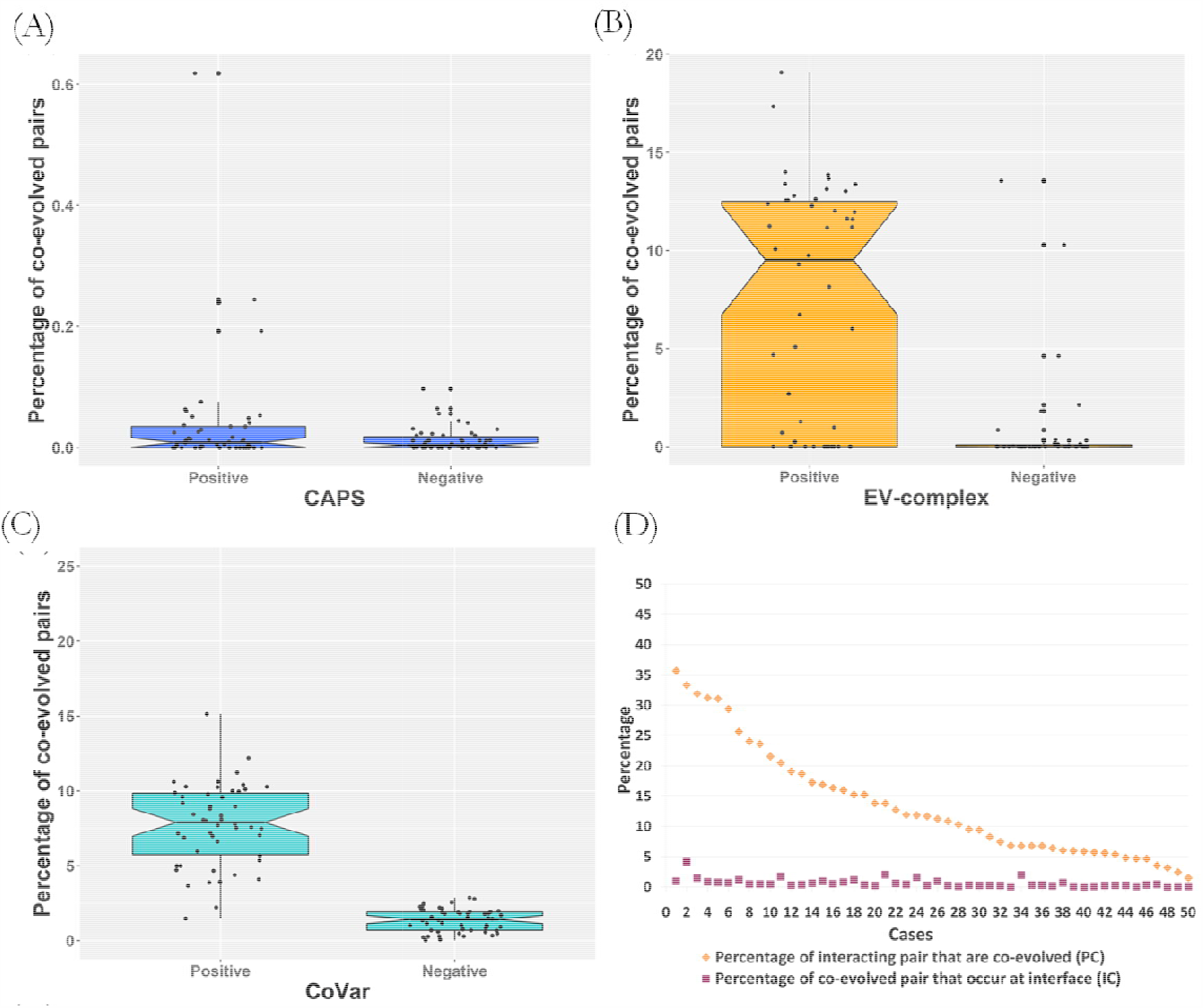
Comparison Co-Var methodology with other inter-protein co-evolution analysis programs. Co-evolved residue positions were mapped onto a reference structure and inter-residue distances were calculated to analyze whether the residues were in close spatial proximity. (A) Distribution of ‘percentage of co-evolved pairs’ predicted by Co-Var in specific inter-residue distance distribution bins (B) Percentage of predicted co-evolved residue pairs that occur at the interface and percentage of interacting pairs that were found to co-evolve among the positive set of complexes analyzed utilizing Co-Var (C) Inter-residue distance distribution for EVcomplex predicted co-evolved pairs (D) Distance distribution analysis for CAPS based prediction of co-evolved pairs in interacting complexes.

The Co-Var methodology devised herein identified that interacting (positive) complexes have ‘percentage of co-evolved pairs’ index in the range of 6-10% while the non-interacting (negative) complexes have much lower co-evolving pairs which lie in the range of 0-2% and as such this index allowed the segregation of the two set of complexes substantially (p-value <= 0.0001) (Figure 4C, Table 1). However, while the distributions of the measures calculated for the positive and negative set of complexes are segregated in these cases as well, significance is lower than that obtained with Co-Var (Figure 3C-D, Table 1). A closer look at the figures also reveals that overlap of ‘percentage of co-evolved pairs’ data points between positive and negative sets are far lesser in Co-Var than that of EV-complex (Figure 4A-C). Therefore, this analysis indicated that the Co-Var methodology performs better to determine whether protein-protein interaction complexes are likely to co-evolve and identifies actual residue pair positions exhibiting such inter-dependent changes. It was identified that a large percentage of co-evolved pairs (nearly 70%) did not lie in close spatial proximity (<10 Å) (Figure 3D). This trend was consistently observed in the co-evolving pairs determined in these complexes with the help of multiple inter-protein co-evolution analysis methodologies (Figure 3B-D).

**Table 1.** Analysis of inter-protein co-evolution utilising Co-Var and other existing methodologies. Statistics for student’s t-test performed utilising co-evolution parameters obtained for interacting (positive) and non-interacting (negative) proteins in Co-Var, MirrorTree, CAPS and EVcomplex has been outlined here.

|  | Co-Var |  | MirrorTree<br>correlation<br>coefficient |  | CAPS |  | EV-complex |  |
| --- | --- | --- | --- | --- | --- | --- | --- | --- |
|  | Positive<br>(n=50) | Negative<br>(n=50) | Positive<br>(n=50) | Negative<br>(n=50) | Positive<br>(n=50) | Negative<br>(n=50) | Positive<br>(n=50) | Negative<br>(n=50) |
| Mean | 7.73 | 1.26 | 0.84 | 0.42 | 0.01 | 0.04 | 7.275 | 0.729 |
| Standard deviation | 2.71 | 0.75 | 0.12 | 0.34 | 0.019 | 0.1 | 6.034 | 2.5 |
| T-test statistics | t=16.2703 df=98 |  | t = 8.2369 df = 98 |  | t=2.0840 df=98 |  | t = 6.8875 df = 96 |  |
| P-value | <= 0.0001 |  | <= 0.0001 |  | 0.0398 |  | <=0.0001 |  |

### Co-evolutionary pairings in intra-protein and inter-protein co-evolution may occur among residue pairs that are not in close spatial proximity

Studies on intra-molecular co-evolution in proteins have suggested that residues in close proximity are likely to be highly co-evolving; alternately distant sites having a functional dependence are likely to co-evolve as well (Fares and Travers 2006, Chakrabarti and Panchenko 2009). Thus, intra-molecular co-evolution was also studied in a set of proteins to study the pattern of inter-residue distances among Co-Var predicted intra-protein co-evolved positions. A similar trend of co-evolving positions occurring in close spatial proximity and in distal regions was observed when we determined co-evolving residues within proteins (252 conserved domain database (CDD) (Marchler-Bauer et al., 2015) protein families) utilising Co-Var, CAPS, MI and PSICOV methods (Figure 5). Additionally, this analysis also suggested that Co-Var may be utilised to study intra-protein co-evolution as well since it predicts a higher percentage of co-evolved pairs that lie in close proximity in comparison to the other existing methodologies. Therefore, considering inter-residue distances among intra-protein co-evolved pairs we found that a higher proportion of pairs occur in close proximity whereas inter-protein co-evolved pairs had a significant fraction of co-evolved pairs that do not lie in close spatial proximity (Figure 3D, Figure 4A). Moreover, in general only about 13.6% (average) of the interface pairs tend to co-evolve in these protein-protein interaction complexes with varying rates among the interaction complexes studied herein (Figure 4D, Supplementary Table 1).

**Figure 5:**
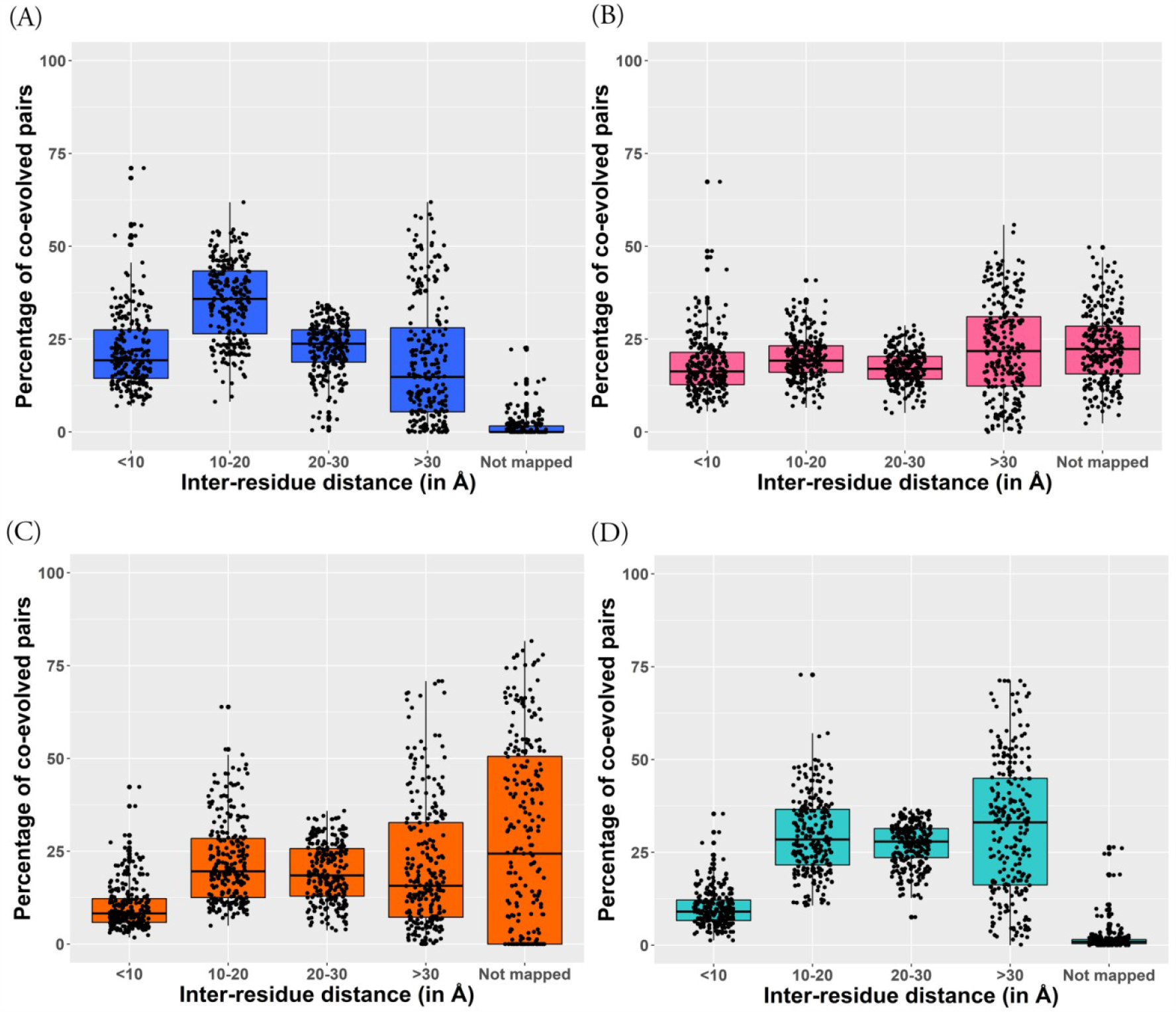
Studying intra-protein co-evolution utilizing Co-Var. Distance distribution analysis of co-evolved positions predicted in intra-protein interaction complexes was considered to evaluate the performance of Co-Var against existing methods for studying intra-protein co-evolution. (A) Inter-residue distance among predicted co-evolved pairs within proteins based on Co-Var. (B) Distance distribution analysis of intra-protein co-evolved pairs according to CAPS. (C) Inter-residue distance among predicted co-evolved pairs within proteins utilizing mutual information. (D) Distance distribution analysis considering intra-protein co-evolving pairs determined in PsiCov.

### Studying co-evolution in hetero-dimeric protein complexes involved in intercellular interactions

Co-evolutionary pairings were determined in hetero-dimeric protein complexes (Table 2) with the help of Co-Var. Predicted co-evolved positions in these selected case studies for protein-protein interaction complexes also suggested that a large fraction of co-evolved positions do not lie at the interface (only about 0.1-0.4% of the total co-evolved pairs lie at the interface) (Table 2). Additionally, in general about 1.5-16% (mean= 5.56%) of the interface pairs were found to co-evolve in the complexes considered and this is in correlation with the observation that transient interfaces exhibit a low degree of co-evolution (Mintseris et al., 2005).

**Table 2:**
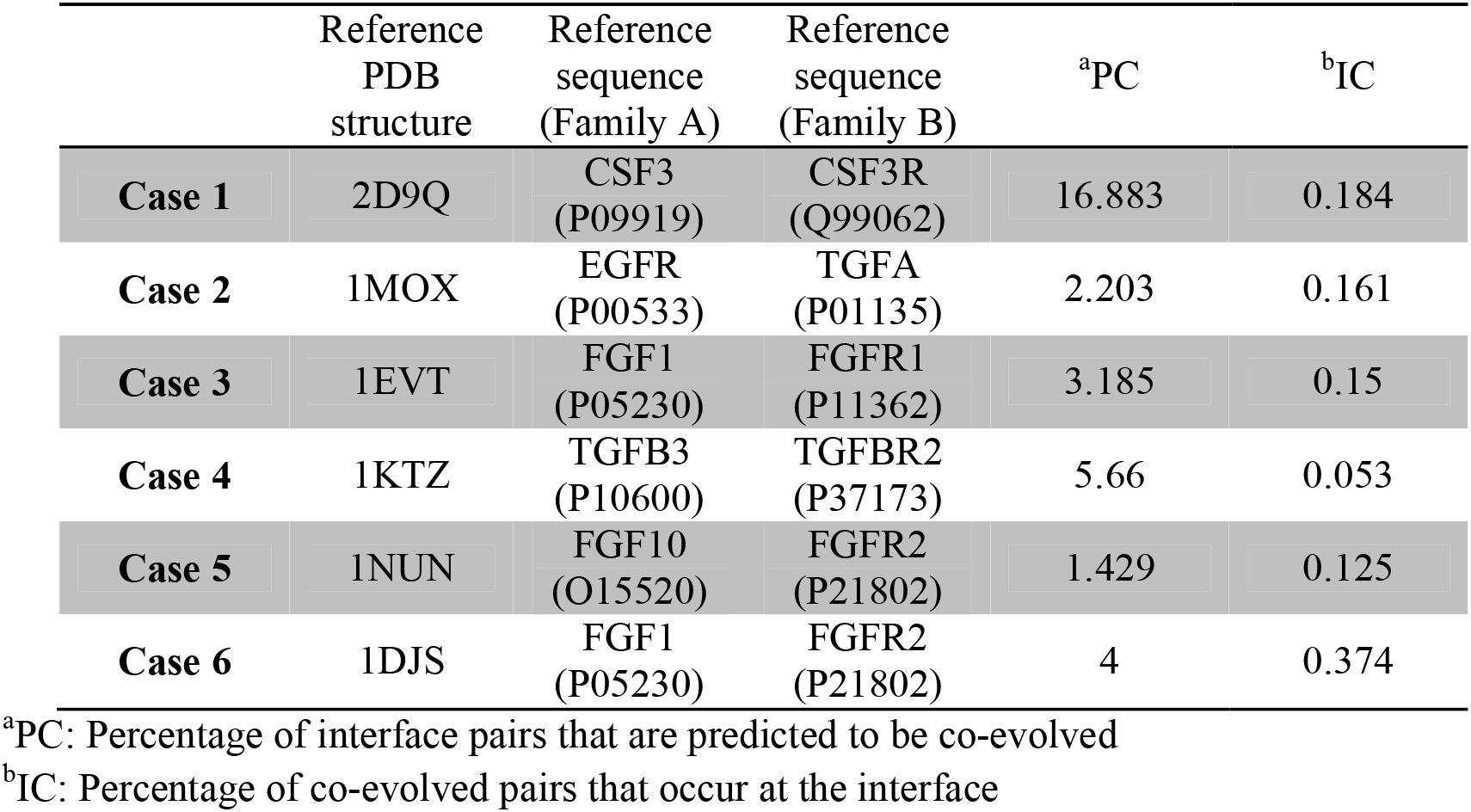
Co-evolution analysis in hetero-dimeric protein complexes involved in inter-cellular interactions. Co-evolutionary pairings were identified utilising Co-Var and a z-score threshold of z<=-1 to study co-evolution in inter-cellular protein interaction complexes.

### Functionally important co-evolutionary pairings occur at interface and non-interface regions in inter-cellular protein interaction complexes

In order to study the probable significance of co-evolutionary pairings which are not occurring in close spatial proximity, we have studied co-evolution patterns in certain receptor-ligand protein hetero-dimers that are known to interact aberrantly during cancer metastasis. A large fraction of co-evolutionary connections between proteins involved in intercellular interactions did not occur in close spatial proximity (Table 2, Supplementary Table 3). However, they were predominantly found to occur only within functional domains of the proteins involved in the interaction (Figure 6). Further, in most of the inter-cellular protein interaction complex case studies analyzed herein about 70-80% of the positions involved in co-evolutionary pairings occur within functional domain regions of the interacting proteins (Table 3). This observation suggests that these residue positions could be biologically relevant for functional integrity of the complex. For instance, in the TGF-A and EGFR interaction complex, co-evolutionary pairings were obtained between 23 out of total 161 residue positions in TGF-A and 251 out of 1210 residue positions in EGFR, respectively. Further, most co-evolutionary pairings were obtained between certain TGF-A (9) and EGFR (91) residues resulting in this tendency to have large number of co-evolutionary connections in the interacting protein partner (Figure 7A). However, co-evolutionary connections among high degree residue positions occur at the interface as well as non-interface regions in this protein pair as well (Figure 7B). In particular, considering the CSF3-CSF3R complex, co-evolutionary pairings were obtained between 68 out of total 207 residue positions in CSF3 and 187 out of 836 residue positions in CSF3R, respectively. However, most co-evolutionary pairings were obtained between certain CSF3 (58) and CSF3R (68) residues only. Here, we observed that certain residue positions exhibit a tendency to have large number of co-evolutionary connections with positions in the interacting protein partner. Thus, these co-evolutionary pairings which exist between residues in CSF3 and CSF3R comprise of 58 residue positions inter-connecting with 68 positions in the interacting protein partner resulting in a large number of co-evolutionary connections among them (Supplementary Figure S1A). Co-evolutionary connections occurring between high degree co-evolved positions in this protein pair are present at the interface as well as non-interface regions and these could be important for the interaction between these proteins (Supplementary Figure S1B). Further, in the TGFB3 and TGFBR2 interaction complex, co-evolutionary pairings were obtained between 104 out of total 412 residue positions in TGFB3, and 95 out of 567 residue positions in TGFBR2 respectively. Further, most co-evolutionary pairings were obtained between certain TGFB3 (53) and TGFBR2 (45) residues again exhibiting this tendency of few residues in one binding partner to have large number of co-evolutionary connections with certain residues in the interacting protein partner (Supplementary Figure S2A). Additionally, co-evolutionary connections were noted among high degree residue positions at the interface and non-interface regions in this protein pair as well (Supplementary Figure S2B). Similarly, the co-evolutionary pairings obtained in the other inter-cellular interaction complexes also exhibit this tendency wherein certain residue positions exhibit a tendency to have larger number of co-evolutionary connections with positions in the interacting protein partner (Supplementary Figures S3,S4,S5). However, these high degree residue positions involved in these co-evolutionary pairings occur in interface (spatially proximal <7Å) and non-interface (spatially distal > 7Å) regions (Figure 8).

**Table 3:** High degree co-evolutionary pairings in inter-protein interaction complexes. Considering residue positions as nodes and co-evolutionary pairings as edges, residue positions that have large number of co-evolutionary connections among them have been determined. Percentage of co-evolved residue positions that have a large number of co-evolutionary connections among them and ones that frequently prone to substitution mutations in cancer are reported.

|  | Protein A | Protein B | <sup>a</sup>CP in FD (%) | <sup>b</sup>HD CP with mutations (%) | <sup>c</sup>Residues forming HD CP (Protein A) | <sup>d</sup>Residues forming HD CP (Protein B) |
| --- | --- | --- | --- | --- | --- | --- |
| **Case1** | CSF3 (P09919) | CSF3R (Q99062) | 78.74 | 45.36 | 58 | 68 |
| **Case 2** | EGFR (P00533) | TGFA (P01135) | 32.74 | 62.78 | 91 | 9 |
| **Case 3** | FGF1 (P05230) | FGFR1 (P11362) | 79.42 | 53.59 | 34 | 23 |
| **Case 4** | TGFB3<br>(P10600) | TGFBR2<br>(P37173) | 84.43 | 55.71 | 53 | 45 |
| **Case 5** | FGF10<br>(O15520) | FGFR2<br>(P21802) | 78.68 | 52.83 | 34 | 31 |
| **Case 6** | FGF1<br>(P05230) | FGFR2<br>(P21802) | 76.68 | 59.40 | 28 | 23 |

<sup>a</sup>**CP in FD (%)**: Percentage of co-evolutionary pairings among residues in functional domains
<sup>b</sup>**HD CP with mutations (%)**: Percentage of high degree co-evolutionary pairings with mutations
<sup>c</sup>**Residues forming HD CP (Protein A)**: Number of residues involved in high degree co-evolutionary pairings in representative protein from family A.
<sup>d</sup>**Residues forming HD CP (Protein B)**: Number of residues involved in high degree co-evolutionary pairings in representative protein from family B.

**Figure 6:**
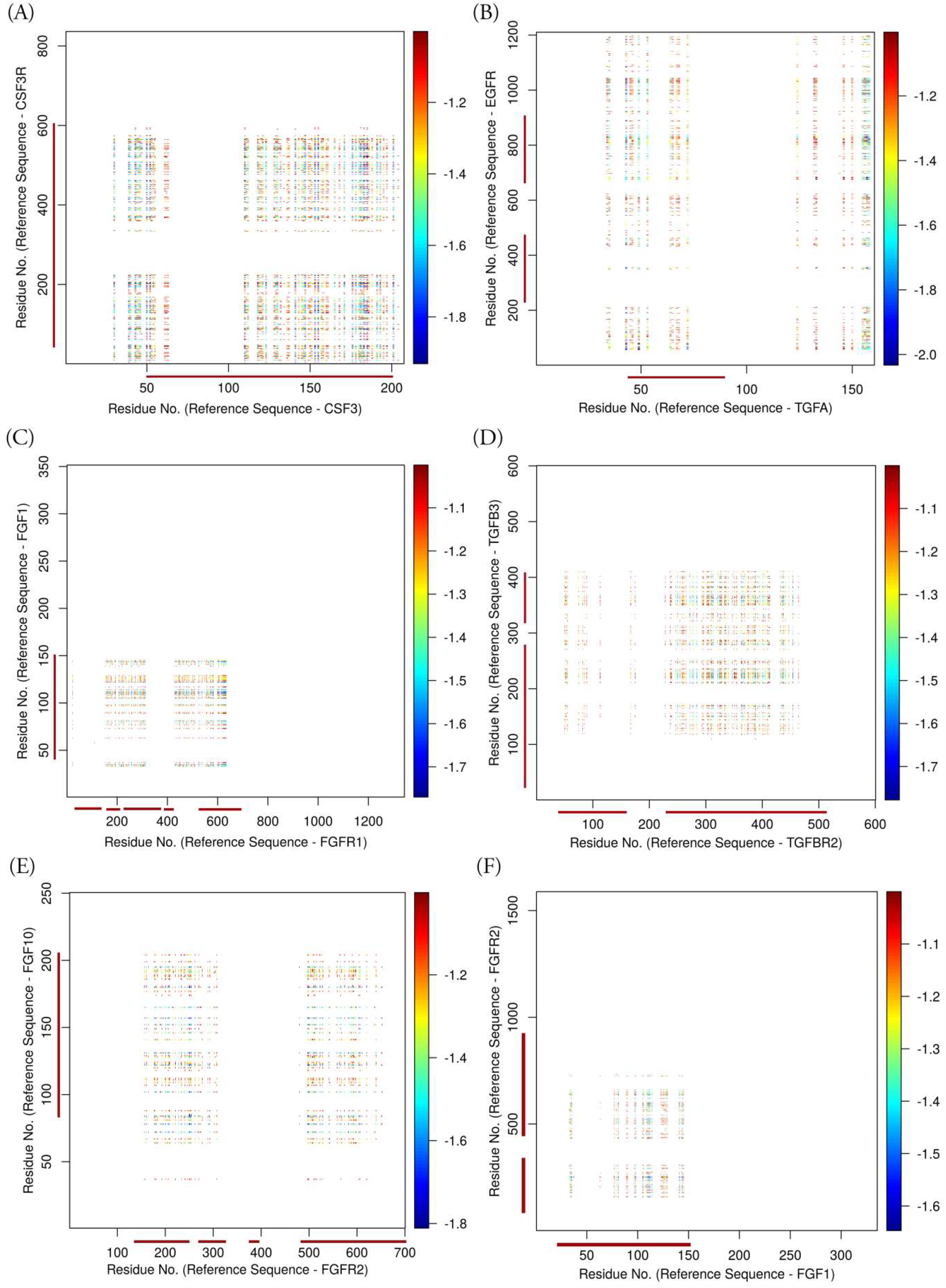
Co-evolving residue positions in inter-protein interactions lie predominantly within functional domain regions within each protein of the complex. Z-scores corresponding to residue positions involved in predicted co-evolutionary pairings (Z-score<=-1) have been plotted (A) Predicted co-evolutionary pairing positions observed between CSF3 and CSF3R.(B) Predicted co-evolutionary pairings positions in TGFA and EGFR complex (C) Predicted co-evolutionary pairing positions occurring between FGF1 and FGFR1 (D) Predicted co-evolutionary pairing positions in TGFBR2 and TGFB3 complex (E) Predicted co-evolutionary pairing positions observed between FGF10 and FGFR2 (F) Predicted co-evolutionary pairing positions in FGF1 and FGFR2 complex.

**Figure 7:**
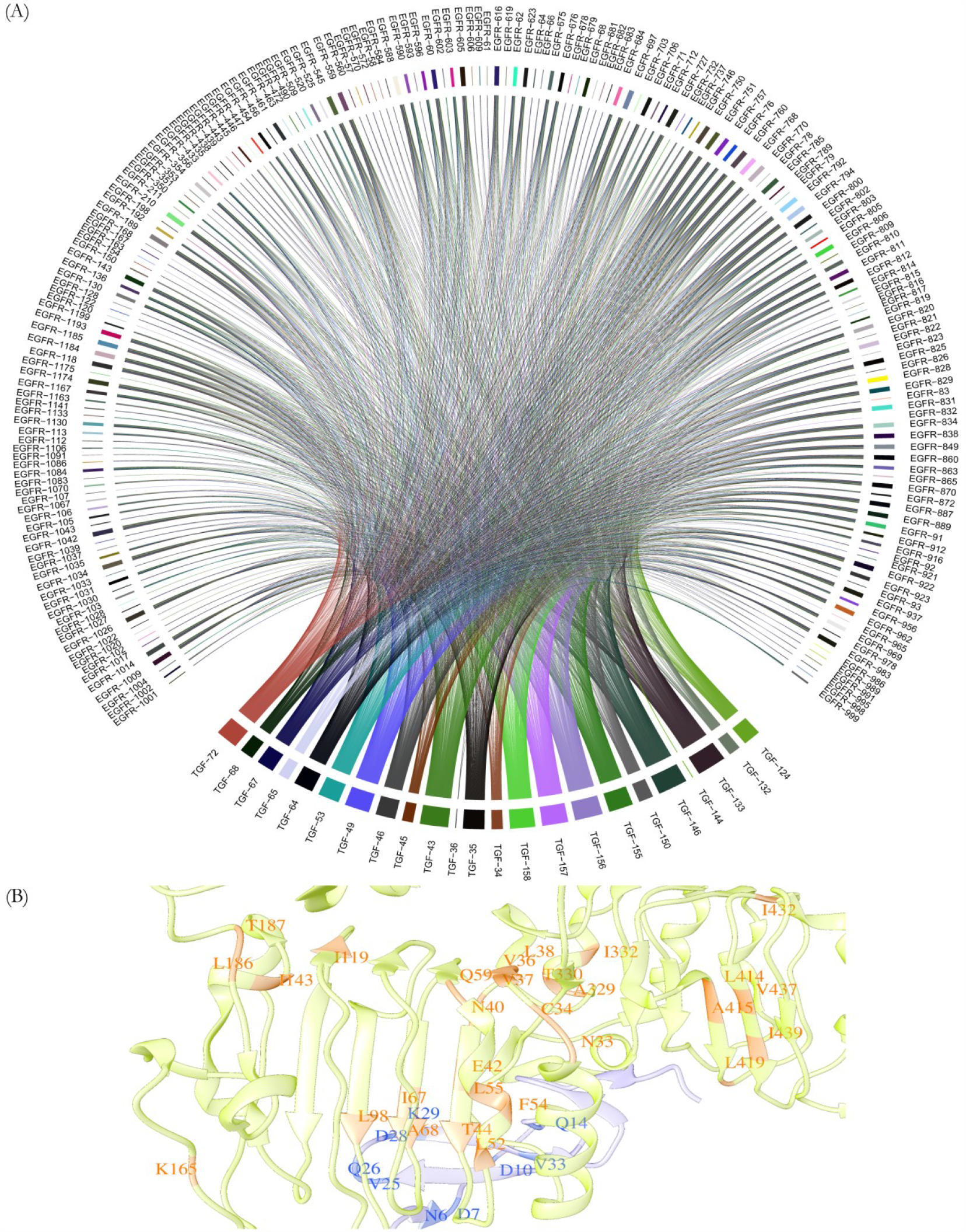
High degree co-evolved positions in inter-cellular protein interaction complex involving TGF-A and EGFR. Residues predicted as co-evolved in inter-protein interaction complexes tend to have a large number of co-evolutionary connections or pairings among them. (A) Co-evolving residues in TGF-A and EGFR that tend to have a large number of co-evolutionary connections (High degree co-evolved positions) or pairings among them are shown here. (B) High degree co-evolved positions mapped onto the reference structure (PDB ID: 1MOX) lie in spatially proximal and distal regions. In the structural representation of co-evolved positions EGFR is depicted in light green while TGFA is depicted is light blue.

**Figure 8:**
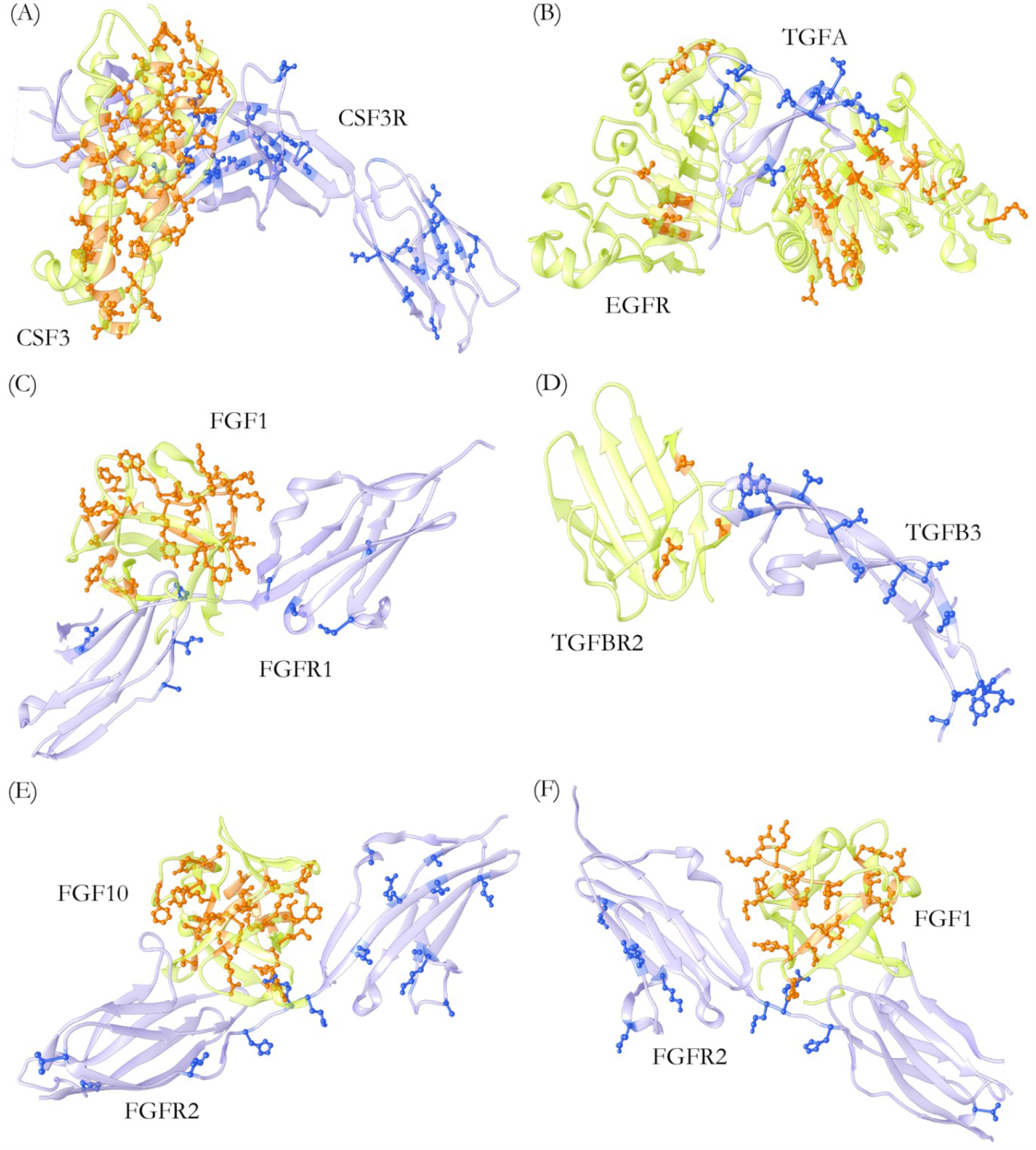
High degree co-evolved positions observed in interface and non-interface regions. Co-evolutionary pairings may include residue positions that have a large number of co-evolutionary connections or pairings among them. Such high degree co-evolved positions when mapped onto reference structures were found to occur in interface and non-interface regions as represented here. High degree co-evolved positions in (A) CSF3 and CSF3R (B) TGFA and EGFR (C) FGF1 and FGFR1 (D) TGFBR2 and TGFB3 (E) FGF10 and FGFR2 (F) FGF1 and FGFR2 complex, respectively are shown in ball and stick models.

### Identification of disease associated changes in intercellular protein-protein interaction complexes involved in cancer metastasis

An interesting observation regarding the high degree co-evolved positions or the residue positions with large number of co-evolutionary connections is that a large fraction among them are frequently prone to substitution mutations prevalent in cancer (Table 3). This finding suggested that residue pairings which are highly co-evolved but frequently show substitution mutations in cancer conditions could potentially be important for conservation of the functional interaction between the interacting proteins. Moreover, we have also studied the residue pairing propensity (frequently observed residue pairs) among the high degree co-evolved positions. It was identified that the residue pairing propensity varies substantially in mutated protein complexes wherein pairs containing amino acids such as glycine, proline, aspartate, glutamate, tryptophan, tyrosine, histidine and glutamine are more frequent (Figure 9). Such substitutions or alterations in pairing propensity at crucial positions that tend to co-evolve are likely to have deleterious functional characteristics in the absence of coordinated compensatory changes. A similar trend is observed in most of the intercellular protein interaction complexes considered herein where the residue pairing propensity among the co-evolved positions gets altered frequently as a result of mutations under diseased conditions (Table 3, Figure 9). Based on these observations it can be postulated that co-evolved residue positions could be frequently mutated in cancers and as such changes at these residue positions may not always be compensatory changes which may result in a perturbed interaction between these proteins. Therefore, with the help of the Co-Var methodology we can predict residue positions in interacting proteins which may or may not be in close proximity but are likely to be functionally relevant or important for an interaction to be maintained between the proteins involved in an intercellular interaction. Further, absence of co-ordinated changes at these at interface and non-interface co-evolving residue positions may lead to disruption of inter-cellular protein-protein interactions and such alterations could be disease associated.

**Figure 9:**
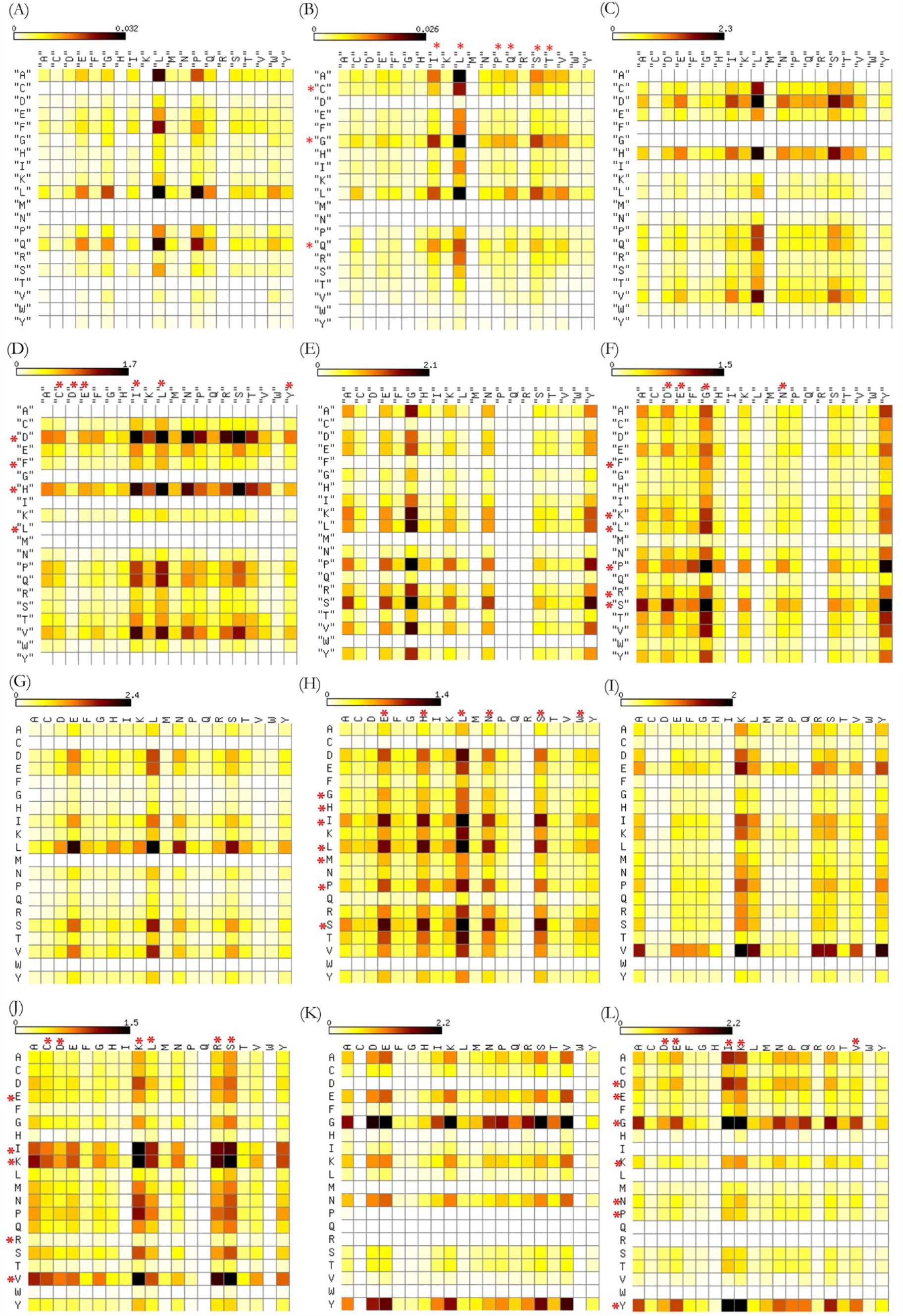
Predicted high degree co-evolved positions may be functionally relevant in protein-protein interactions. Residue pairing propensity at the high degree co-evolved positions in reference protein sequences of the complex and the altered pairing propensity based on the observed substitution mutations have been compared. High degree co-evolved positions are frequently prone to substitution mutations and altered residue pairing propensity at these positions have been highlighted with *. (A) Pairing propensity in native CSF3 and CSF3R complex (B) Altered pairing propensity in CSF3 and CSF3R complex (C) Pairing propensity in native TGFA and EGFR complex (D) Altered pairing propensity in mutated TGFA and EGFR complex (E) Pairing propensity in native FGF1 and FGFR1 complex (F) Altered pairing propensity in mutated FGF1 and FGFR1 complex (G) Pairing propensity in native TGFBR2 and TGFB3 complex (H) Altered pairing propensity in mutated TGFBR2 and TGFB3 complex (I) Pairing propensity in native FGF10 and FGFR2 complex (J) Altered pairing propensity in mutated FGF10 and FGFR2 complex (K) Pairing propensity in native FGF1 and FGFR2 complex (L) Altered pairing propensity in mutated FGF1 and FGFR2 complex.

Moreover, intra-molecular co-evolution analysis of the complex constituent proteins demonstrated that positions important for protein stability or function are also involved in forming extensive high degree co-evolutionary pairings in inter-protein interaction complexes as well. Therefore, residue positions important for intra-protein stability or function may additionally influence inter-protein co-evolution interactions as well (Table 4).

**Table 4:**
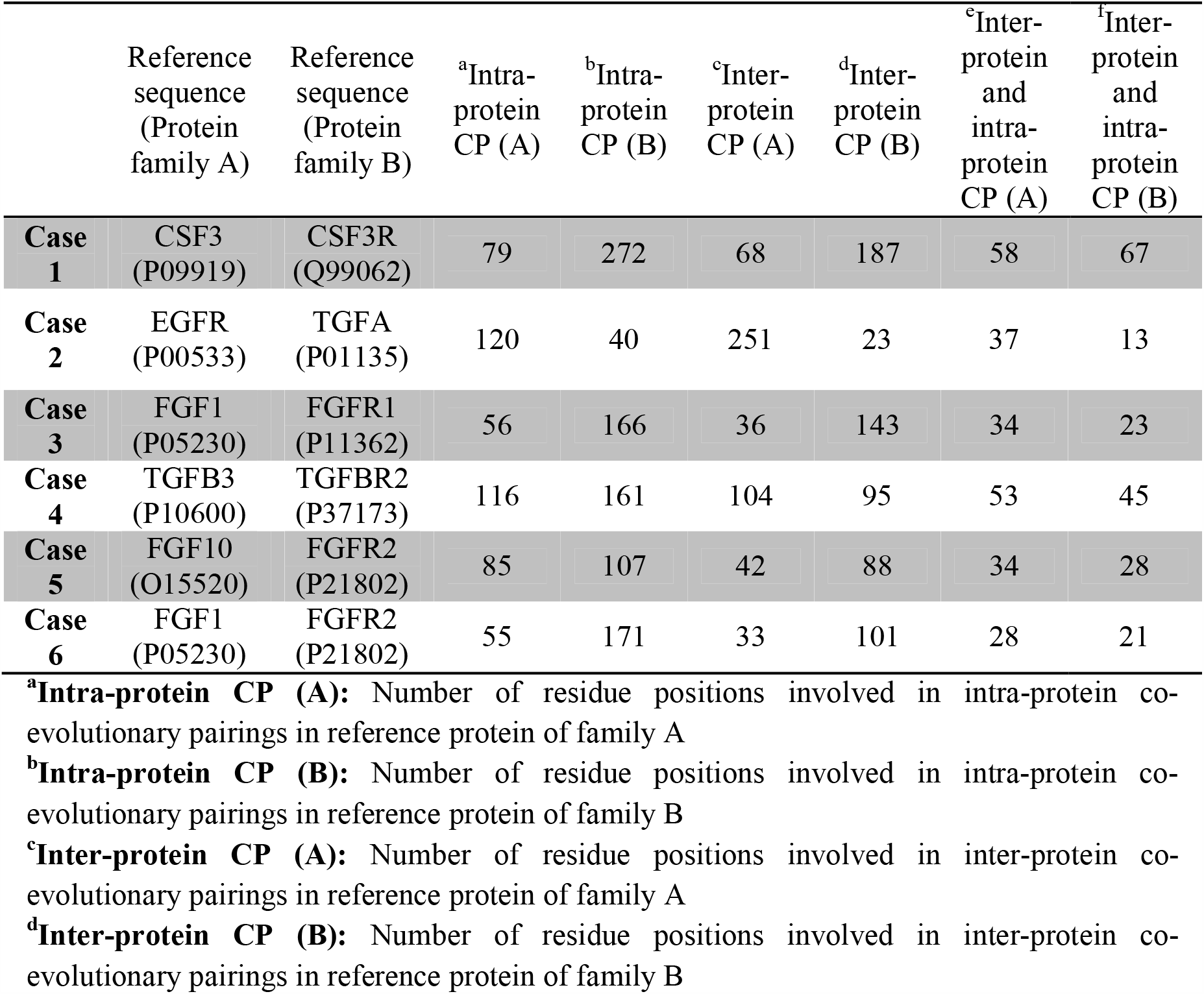
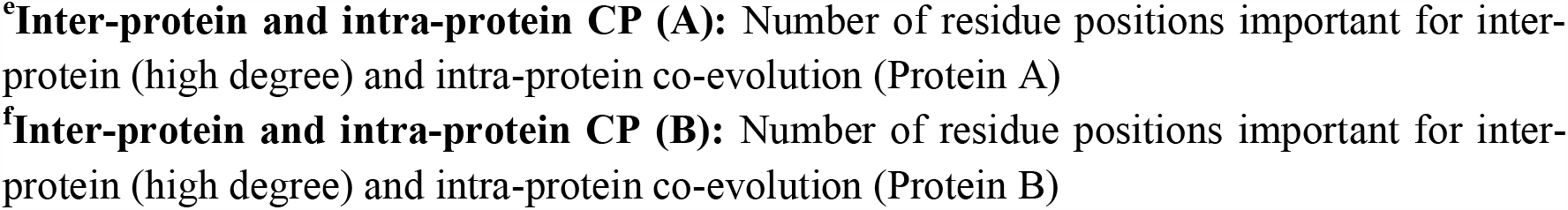
Intra-protein co-evolving positions in proteins constituting a complex are also predicted as high degree inter-protein co-evolved positions. Inter-protein co-evolving residue positions that have large number of co-evolutionary connections (high degree) between proteins are likely to be important for intra-molecular co-evolution as well.

|  | Reference<br>sequence<br>(Protein<br>family A) | Reference<br>sequence<br>(Protein<br>family B) | <sup>a</sup>Intra-protein<br>CP (A) | <sup>b</sup>Intra-protein<br>CP (B) | <sup>c</sup>Inter-protein<br>CP (A) | <sup>d</sup>Inter-protein<br>CP (B) | <sup>c</sup>Inter-protein<br>and<br>intra-protein<br>CP (A) | <sup>f</sup>Inter-protein<br>and<br>intra-protein<br>CP (B) |
| --- | --- | --- | --- | --- | --- | --- | --- | --- |
| **Case 1** | CSF3<br>(P09919) | CSF3R<br>(Q99062) | 79 | 272 | 68 | 187 | 58 | 67 |
| **Case 2** | EGFR<br>(P00533) | TGFA<br>(P01135) | 120 | 40 | 251 | 23 | 37 | 13 |
| **Case 3** | FGF1<br>(P05230) | FGFR1<br>(P11362) | 56 | 166 | 36 | 143 | 34 | 23 |
| **Case 4** | TGFB3<br>(P10600) | TGFBR2<br>(P37173) | 116 | 161 | 104 | 95 | 53 | 45 |
| **Case 5** | FGF10<br>(O15520) | FGFR2<br>(P21802) | 85 | 107 | 42 | 88 | 34 | 28 |
| **Case 6** | FGF1<br>(P05230) | FGFR2<br>(P21802) | 55 | 171 | 33 | 101 | 28 | 21 |

<sup>a</sup>**Intra-protein CP (A)**: Number of residue positions involved in intra-protein co-evolutionary pairings in reference protein of family A
<sup>b</sup>**Intra-protein CP (B)**: Number of residue positions involved in intra-protein co-evolutionary pairings in reference protein of family B
<sup>c</sup>**Inter-protein CP (A)**: Number of residue positions involved in inter-protein co-evolutionary pairings in reference protein of family A
<sup>d</sup>**Inter-protein CP (B)**: Number of residue positions involved in inter-protein co-evolutionary pairings in reference protein of family B
**<sup>e</sup>Inter-protein and intra-protein CP (A):** Number of residue positions important for inter-protein (high degree) and intra-protein co-evolution (Protein A)
**<sup>f</sup>Inter-protein and intra-protein CP (B):** Number of residue positions important for inter-protein (high degree) and intra-protein co-evolution (Protein B)

### Co-Var web server for studying intra-protein and inter-molecular co-evolution

A web server for analyzing intra-protein and inter-molecular co-evolution is available online at http://www.hpppi.iicb.res.in/ishi/covar/index.html (Figure 10A). During the inter-protein co-evolution analysis, co-evolutionary pairings are determined based on the Co-Var methodology and reference sequence mapped co-evolving positions are reported with the help of a surface plot representation. The Co-Var score and Z-score for threshold selected co-evolutionary pairings are depicted (Figure 10B). High degree co-evolved positions and/or co-evolved positions in close spatial proximity are displayed on the reference structure provided and the list of co-evolving positions in close spatial proximity may be downloaded (Figure 10B). Further, a distance distribution plot of the inter-residue distances among co-evolved position pairs provided can be utilized to get an idea about whether co-evolutionary pairings are occurring in close proximity or among spatially distant residue positions. Additionally, high degree co-evolved positions that are likely to be important for maintaining inter-protein functional interaction are also determined and the same are provided as lists. The final results of the analysis are mailed to the e-mail address provided and are easily available for download.

**Figure 10:**
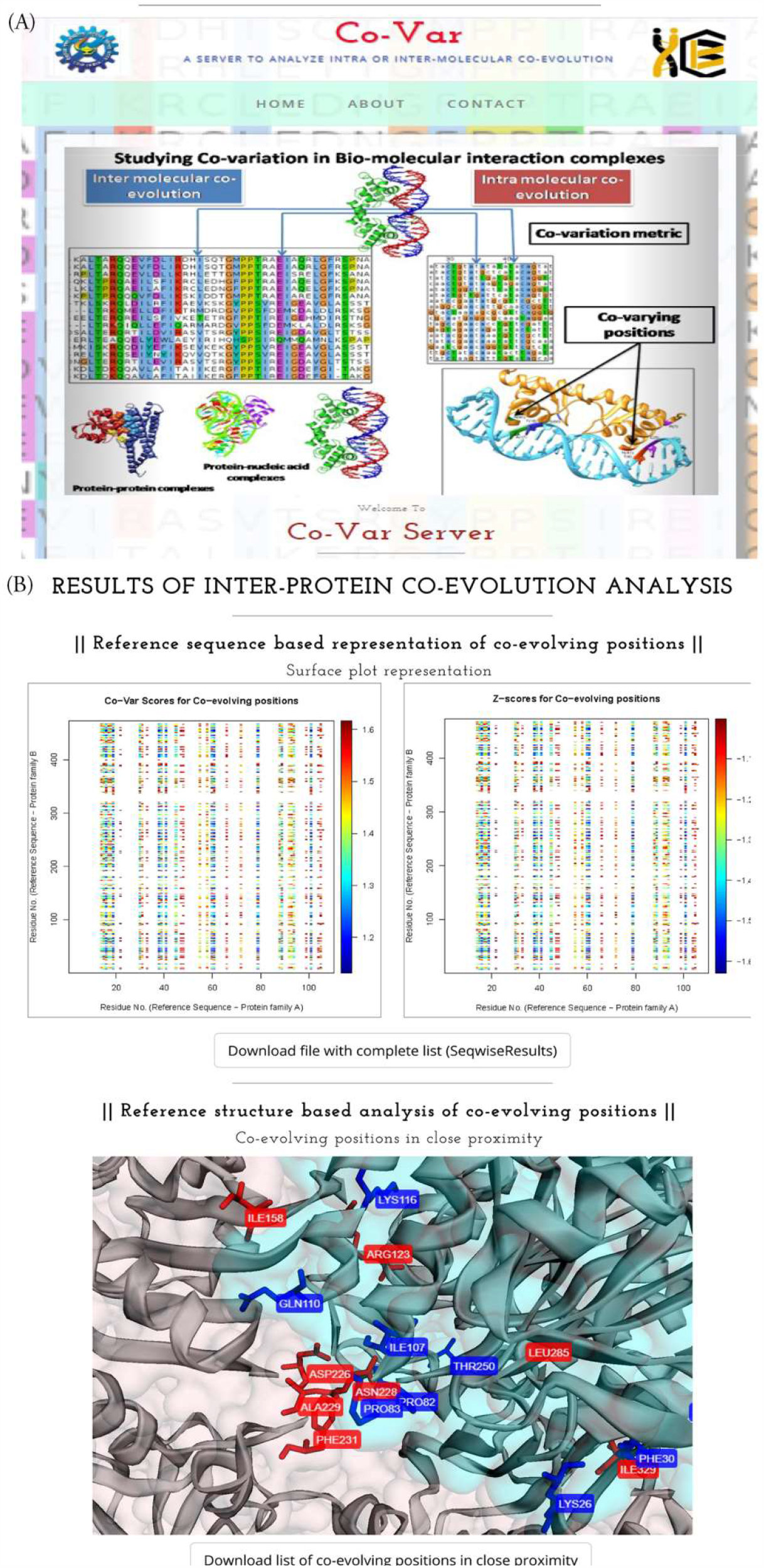
Co-Var web server to study intra-protein and inter-molecular co-evolution. (A) Co-Var web-server user interface (B) Snap-shot of inter-protein co-evolution analysis results provided by Co-Var web-server.

## Discussion

Molecular co-evolution refers to a phenomenon where a change in one locus is likely to affect the selection pressure at another locus and a reciprocal change may occur reflecting a direct evolutionary interaction. Such an evolutionary interaction could be occurring between sites within a single protein referred to as intra-protein co-evolution or between different proteins in which case it is referred to as inter-protein co-evolution (Lovell and Robertson, 2010). In this work, we have developed the Co-Var methodology which utilizes mutual information and Bhattacharyya co-efficient to study intra-protein and inter-molecular co-evolution. Multiple methodologies have previously been developed to study intra-protein (MI, CAPS, and PSICOV) and inter-protein (CAPS, EV-complex) co-evolution (Fares and Travers 2006; Korber et al., 1993; Jones et al., 2012; Hopf et al., 2014). However, our approach has a number of advantages such as non-dependence on identical alignment lengths and a reduced influence of phylogenetic relationships of the organisms/species represented in the alignment. This is because supplied sequences in the alignment are randomly selected for the analysis and alignment shuffling is also performed. Further, the Co-Var methodology described here has also been implemented into an easy to use more generic Co-Var web server platform. Herein, probable co-evolutionary connections within proteins and across biomolecules or complexes, such as protein-protein, can be estimated and their structural and functional relevance can be judged. Moreover, the inter-protein co-evolution analysis platform has been extensively validated and certain case scenarios have been studied in detail to identify whether co-evolutionary couplings occur at the interaction interface or other regions important for complex recognition and formation.

Studies pertaining to intra-molecular co-evolution have identified that coevolving positions occur in two different categories. The first set comprises of positions that coevolve with only one or two other positions and often exhibit direct amino acid side-chain interactions with their coevolving partner in close proximity. However, the second set includes positions that coevolve with many other positions which are predominantly located in regions critical for protein function, for instance active sites or regions involved in intermolecular interactions and recognition (Gloor et al, 2005; Chakrabarti and Panchenko 2009). In a similar manner the Co-Var methodology can be utilised to study intra-molecular co-evolution in proteins. For intra-molecular co-evolution analysis using the CDD protein families we found that in general it predicts a higher fraction of co-evolved residues in close proximity than the other programs that have been considered during this analysis. Co-evolution is also evident in biological systems where the interaction patterns have to be maintained while the interactions continue to evolve and acquire new functions and/or avoid crosstalk with other available systems. This scenario is prevalent in signalling cascades, where a rapid divergence may occur to avoid interference with the original system. However, such change is generally compensated by the interacting partners so as to maintain a functional cascade (Ochoa and Pazos, 2014). Inter-molecular coevolution studies in proteins have shown that residues in close spatial proximity at the interaction interface generally exhibit a higher tendency to co-evolve than other residue pairs predicted as co-evolving which are spatially separated (Mintseris and Weng, 2005; Anishchenko et al., 2017). Here, in our analysis, we could find that a certain fraction of co-evolving residue pairs are predicted in spatially separated positions via multiple inter-protein co-evolution analysis programs that we have utilized. Further, these co-evolutionary pairings that occur at non-interface regions occur among residues present in functional domains or residues that have roles in intra-protein co-evolution in individual proteins involved in the complex. Moreover, with the help of inter-protein co-evolution analysis of intercellular complexes involved in cancer metastasis we could find that certain receptor positions share many co-evolutionary pairing connections with certain ligand positions only. Additionally, these high degree co-evolving positions have been found to be frequently prone to mis-sense substitution mutations in cancer and as such absence of co-ordinated changes at these positions may contribute to an altered interaction in these complexes. Therefore, the Co-Var methodology allows one to predict co-evolving residue pair positions; alterations at which could be functionally detrimental for a protein-protein interaction to occur. Moreover, it has been identified that co-evolutionary pairings crucial for functional interactions in inter-protein complexes may occur in close spatial proximity or at non-interface regions. These residue positions involved in these co-evolutionary pairings are frequently prone to substitution mutations in cancer and occur in functional domains or are important for intra-molecular co-evolution within the proteins involved in the complex. Thus, the information theory based Co-Var measure may be utilised to study interacting proteins that are co-evolving and to determine co-evolutionary pairings among residues that could be structurally or functionally relevant for inter-protein interactions in particular.

## Acknowledgments

Authors would like to acknowledge the initial contribution of Abhijit Chakraborty in the project. We would also like to thank Sunandan Dhar for his involvement in this work during his summer project tenure. SC acknowledges CSIR-Indian Institute of Chemical Biology (IICB) for infrastructural support. IM is thankful to CSIR for her fellowship.

## Author’s contributions

SC and IM formulated the study design. IM performed the experiments. SC and IM analyzed and interpreted the results. The manuscript was prepared by IM and SC. All authors read and approved the final version of the manuscript.

## Data Availability

The data pertaining to the conclusions in this article are available in the article and in its online supplementary material. Data utilized for arriving at the conclusions presented in the work is available upon request. A local version of the Co-Var method to determine co-evolutionary pairings in biomolecules is available for download from the Co-Var web server (http://www.hpppi.iicb.res.in/ishi/covar/download/covar-loc.zip) and/or the GitHub repository (https://github.com/Ishita2690/Co-Var).

