## Supplementary Information for "Co-evolutionary Landscape at the Interface and Non-Interface Regions of Protein-Protein Interaction Complexes"

**Figure S1: High degree co-evolved positions in inter-cellular protein interaction complexes. A.** Co-evolving residues in CSF3 (GCSF) and CSF3R (GCSFR) that tend to have a large number of co-evolutionary connections (high degree co-evolved positions) or pairings among them are shown here. **B.** High degree co-evolved positions mapped onto the reference structure (PDB ID: 2D9Q) lie in spatially proximal and distal regions. In the structural representation of co-evolved positions CSF3 is depicted in light green while CSF3R is depicted is light blue.

(A)

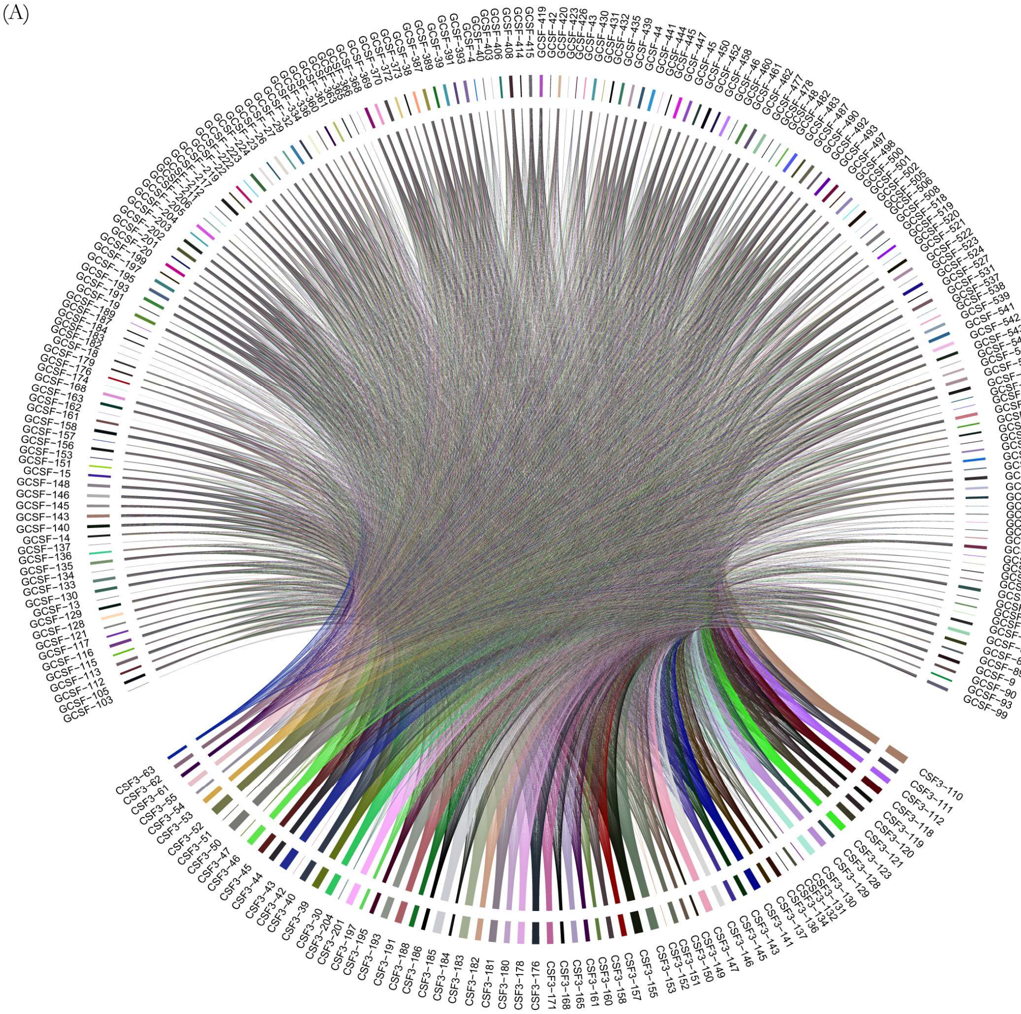

(B)

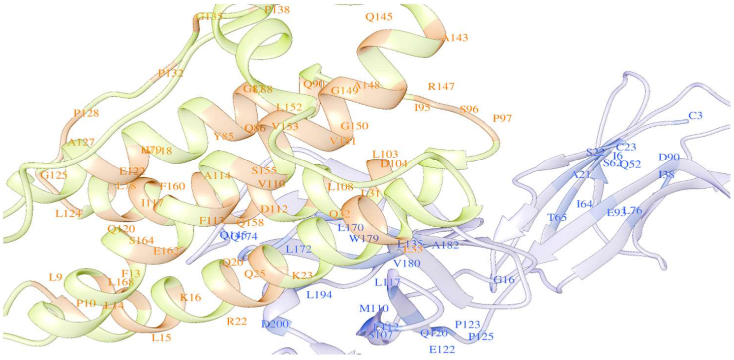

**Figure S2: Predicted high degree co-evolved positions in inter-protein interaction complex between TGFB3 and TGFBR2**

**A.** Co-evolving residues in TGFB3 and TGFBR2 that tend to have a large number of co-evolutionary connections (High degree co-evolved positions) or pairings among them are shown here. **B.** High degree co-evolved positions mapped onto the reference structure (PDB ID: 1KTZ) lie in spatially proximal and distal regions. In the structural representation of co-evolved positions TGFBR2 is depicted in light green while TGFB3 is depicted is light blue.

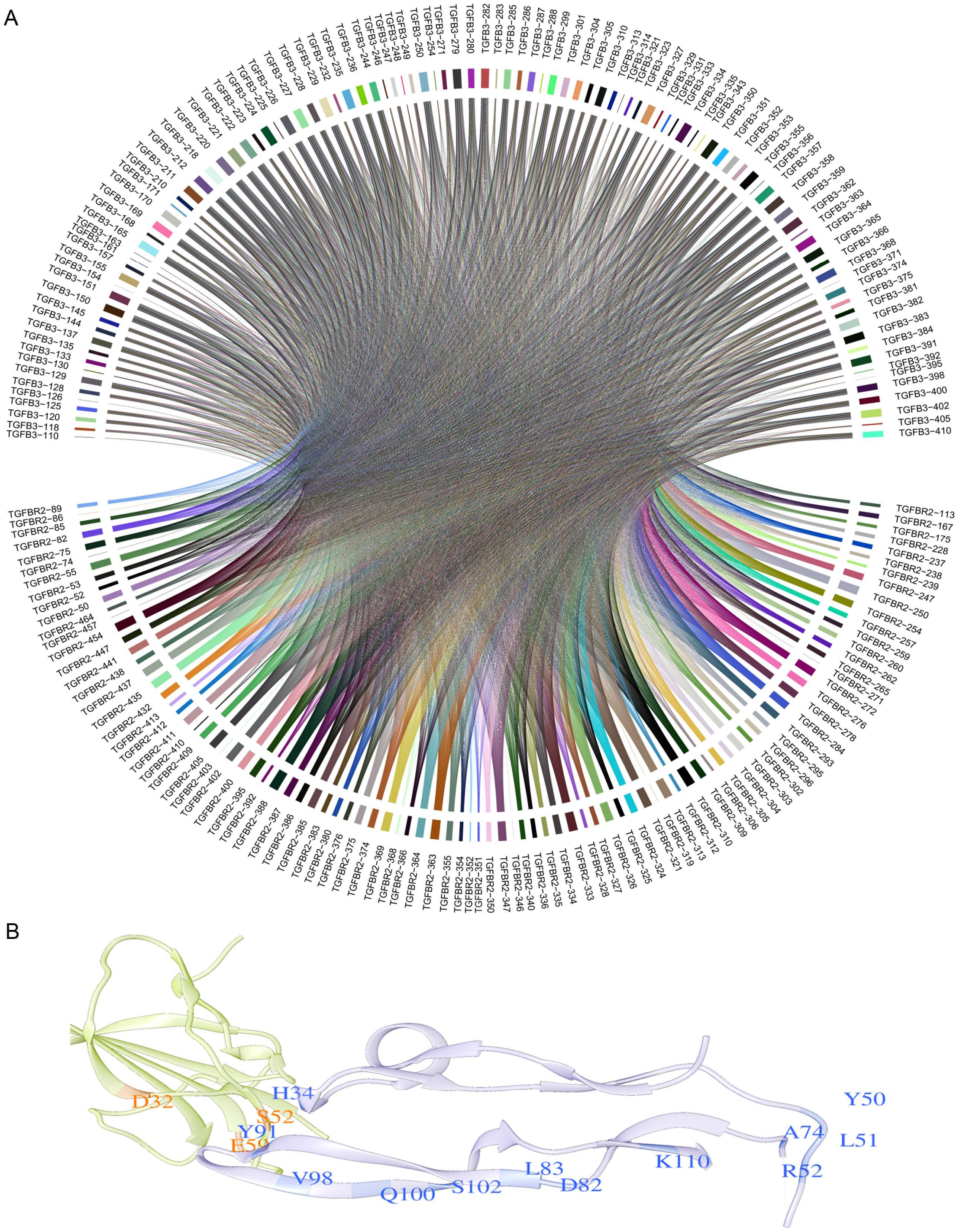

**Figure S3: Residues predicted as co-evolved in FGF1-FGFR1 complex tend to have a large number of co-evolutionary connections or pairings among them A.** Co-evolving residues in FGF1 and FGFR1 that tend to have a large number of co-evolutionary connections (High degree co-evolved positions) or pairings among them are shown here. **B.** High degree co-evolved positions mapped onto the reference structure (PDB ID: 1EVT) lie in spatially proximal and distal regions. In the structural representation of co-evolved positions FGF1 is depicted in light green while FGFR1 is depicted is light blue.

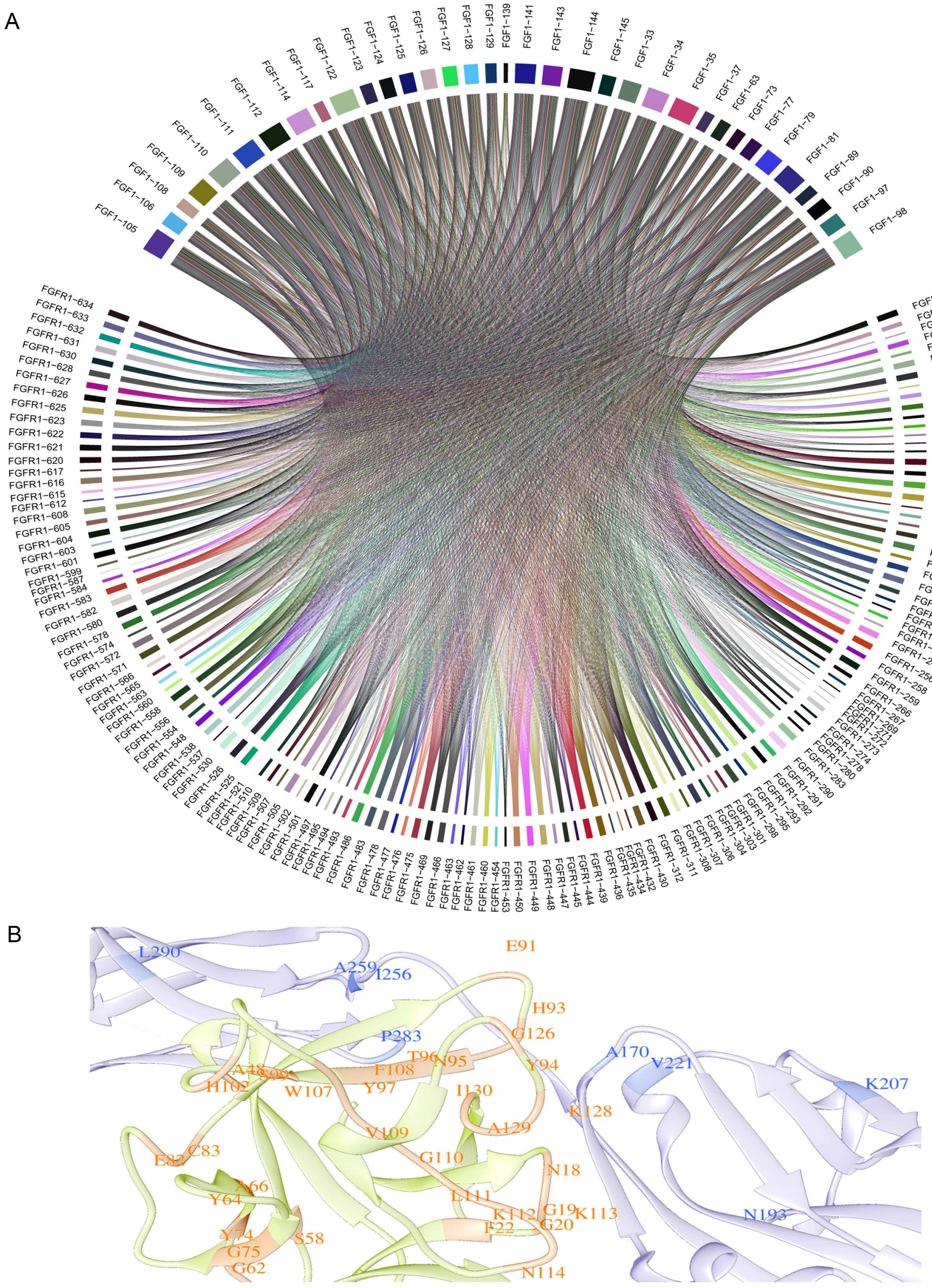

**Figure S4: Inter-cellular protein interaction complex involving FGF10 and FGFR2 have a large number high degree co-evolved positions**

**A**. Co-evolving residues in FGF10 and FGFR2 that tend to have a large number of co-evolutionary connections (High degree co-evolved positions) or pairings among them are shown here. **B.** High degree co-evolved positions mapped onto the reference structure (PDB ID: 1NUN) lie in spatially proximal and distal regions. In the structural representation of co-evolved positions FGF10 is depicted in light green while FGFR2 is depicted is light blue.

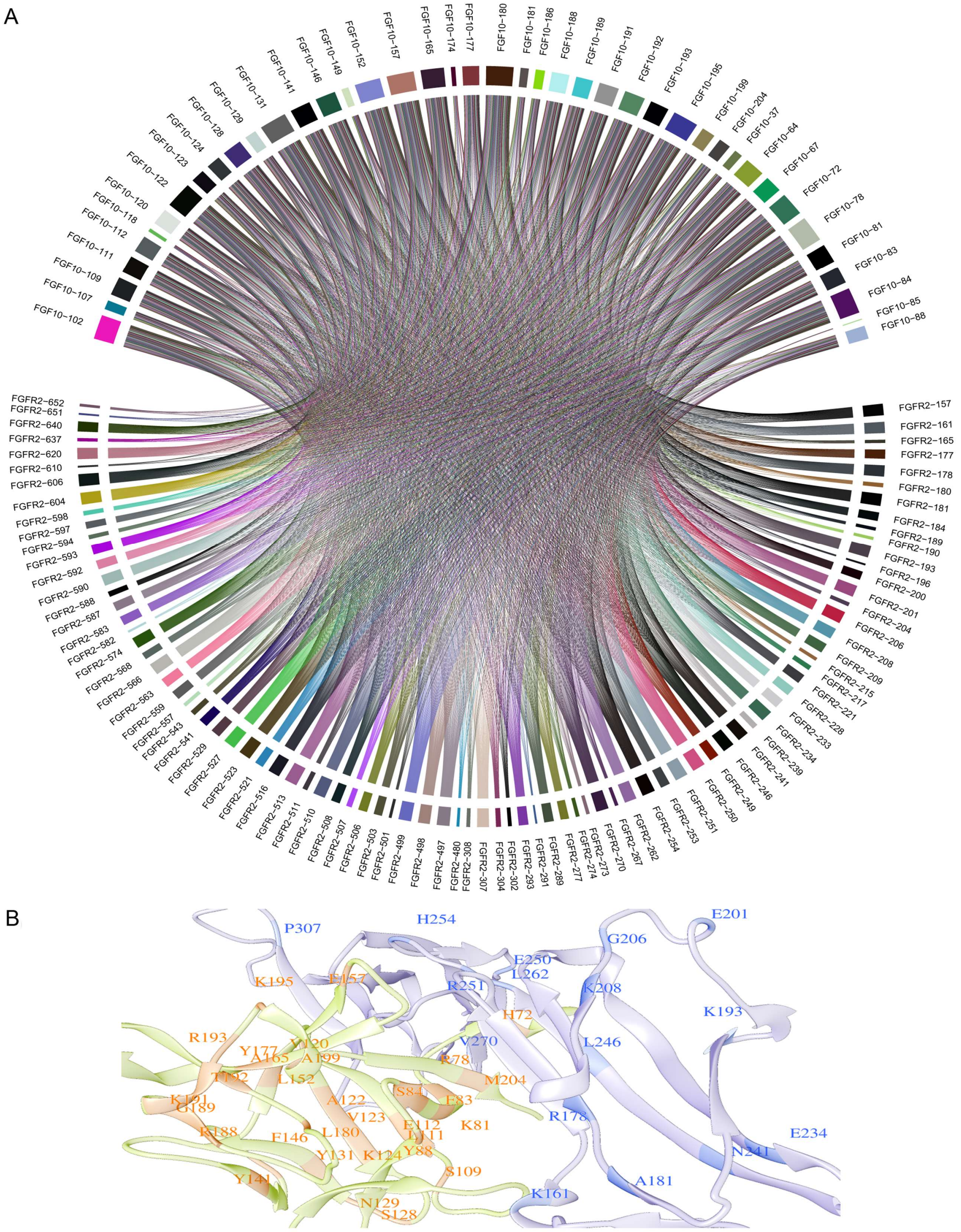

**Figure S5: High degree co-evolved positions in FGF1-FGFR2 complex**

**A**. Co-evolving residues in FGF1 and FGFR2 that tend to have a large number of co-evolutionary connections (High degree co-evolved positions) or pairings among them are shown here. **B.** High degree co-evolved positions mapped onto the reference structure (PDB ID: 1DJS) lie in spatially proximal and distal regions. In the structural representation of co-evolved positions FGF1 is depicted in light green while FGFR2 is depicted is light blue.

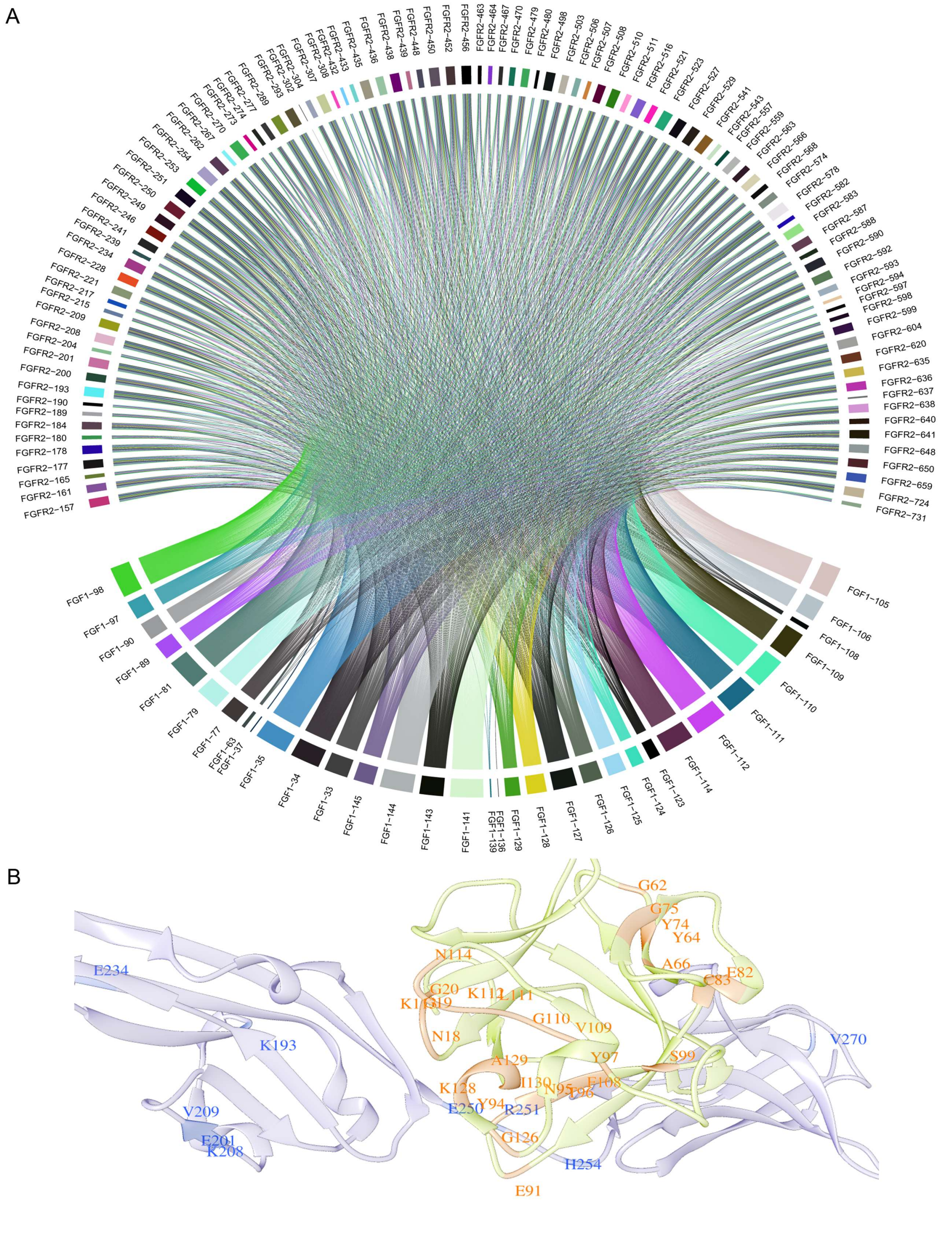

**Table S1: Determining the applicability of Co-Var in studying inter-protein co-evolution analysis**

Details about interacting proteins (Positive set) and non-interacting proteins (Negative set) considered for the protein-protein co-evolution validation and statistics for their analysis in Co-Var, MirrorTree, CAPS and EV-complex is listed here (attached .xls).

**Table S2: Evaluating the performance of Co-Var measure in studying intra-protein co-evolution**

List of proteins considered for the intra-protein co-evolution analysis programs (Co-Var, CAPS, Mutual information and PsiCov) and the details for the same are outlined here (attached .xls).

**Table S3: Co-evolutionary pairings in inter-cellular protein interaction complexes lie in interface and non-interface regions**

| | Reference PDB structure | Reference sequence (Protein family A) | Reference sequence (Protein family B) | Percentage of co-evolved pairs having respective inter-residue distances | | | |
|---|---|---|---|---|---|---|---|
| | | | | <=10Å | 10-20Å | 20-30Å | >30Å |
| **Case 1** | 2D9Q | CSF3 (P09919) | CSF3R (Q99062) | 1.40 | 9.32 | 18.35 | 70.93 |
| **Case 2** | 1MOX | EGFR (P00533) | TGFA (P01135) | 3.83 | 17.02 | 37.02 | 42.13 |
| **Case 3** | 1EVT | FGF1 (P05230) | FGFR1 (P11362) | 2.42 | 17.55 | 33.28 | 46.76 |
| **Case 4** | 1KTZ | TGFB3 (P10600) | TGFBR2 (P37173) | 7.75 | 14.73 | 17.83 | 59.69 |
| **Case 5** | 1NUN | FGF10 (O15520) | FGFR2 (P21802) | 1.83 | 15.49 | 33.42 | 49.26 |
| **Case 6** | 1DJS | FGF1 (P05230) | FGFR2 (P21802) | 3.73 | 17.57 | 33.45 | 45.25 |
